# The building blocks of social structure: simulating constraints of social network analysis for inference about group-level properties

**DOI:** 10.64898/2026.09.17.752303

**Authors:** James Brooks, Gal Badihi, Liran Samuni

## Abstract

1. The interdisciplinary usage of Social Network Analysis (SNA) means that, when researchers calculate global network metrics, the research interest can range from the ultimate ecological and evolutionary pressures that select for specific group-level structures or the proximate mechanisms by which those structures emerge. Despite major methodological advances, social structures measured via SNA metrics are rarely connected to the underlying forces shaped by latent individual dispositions, ecological constraints, and their interplay with group-level affordances.
2. We build a model to simulate fission-fusion spatial association data from a specified set of parameters in order to address the ‘black box’ of animal SNA. We broke down animal social systems into three building blocks: individual dispositions, external factors, and the social structure that emerges. Each of these building blocks was represented by one or more model parameters, which together describe a set of underlying processes that lead to different measurable features of animal social structure, which we quantified with commonly used global network metrics within an SNA framework (e.g., clustering coefficient, density, modularity).
3. Holding all other parameters constant, we find that subtle changes to 1) demography (i.e., group size) and ecology (i.e., size and variability of foraging parties), 2) variation of individual behaviour and related observation bias, 3) imposed group sub-structures, and 4) observation effort, affect all network metrics calculated non-linearly and in some cases non-monotonically.
4. Our findings indicate that failure to account for the underlying behavioural processes can bias inference drawn from SNA. This model and conceptual framework provide a novel tool with which to begin opening the black box of SNA and to systematically address the evolution of diverse group-level social structures.

## 1. Introduction

Social systems have evolved as a balance of costs and benefits, tailored to the ecological and social challenges faced by individuals. Far from being a static trait, individuals both adapt to and reshape their social environment, giving rise to dynamic social phenotypes that reflect ongoing feedback between individual behavior and relationships, group demography, culture, and the group’s broader socio-ecology (Siracusa et al., 2026; Sueur et al., 2026). How such feedback processes between individuals and their wider socio-ecology shape the ways in which societies are organized and structured is a crucial question at the core of human and nonhuman social evolution.

Attempting to understand and capture social variation entails recognizing that social preferences, relationships, and structures cannot be directly observed, but must be inferred from patterns of behaviour (Hinde, 1976). Scholars have increasingly turned towards formalization and quantification of the layers of Hinde’s model. For example, the latent layers framework (De Moor, Hart, et al., 2025) emphasizes the need to distinguish the *observed* network–individual and social behaviour that we record–from latent *realized* networks that we build using these observations and the underlying social preferences (*preference* network). While group-level properties (i.e., the *realized* network) reflect and shape patterns and distributions of dyadic relationships (i.e., the *preference* network), both cannot be observed directly but must be inferred from observation data (i.e., the *observed* network). This parallels concepts in Kawam et al. (2025), who distinguish theoretical constructs such as relationships and preferences, the interactions that reflect them, and the incomplete observational data used to infer these constructs. Relatedly, Neumann and Fischer (2026) demonstrate the importance of, and develop tools for, decomposing the processes of individual gregariousness and dyadic affinity in shaping social interaction data. These works provide a valuable conceptual and methodological toolbox for revealing and disentangling underlying behavioural dispositions from latent group-level structures. Yet, they also highlight the complexity of social systems and the multiple layers at which biases and distortions can arise—particularly in comparative contexts where systems differ in many respects simultaneously. Breaking down latent social networks into the underlying dyadic preferences and interactions is critical for understanding the proximate behavioural mechanisms that shape group-level structures. Substantially less conceptual investment has been dedicated to unpacking how ultimate evolutionary and ecological pressures act on group-level structure, and how these structures feedback to shape individual preferences and behaviour.

In addition to understanding how latent social networks and individual dispositions and preferences drive observable behaviour, studies of social systems must also ensure that empirical measures reflect *true* variation in social phenotypes rather than variation driven by stochasticity, observational biases, and confounding factors (Mielke et al., 2026). For example, demographic factors such as group size and composition can directly shape social systems in biologically meaningful ways (e.g., by constraining partner choice or altering competition), but they also affect how variation in these systems is captured by commonly used metrics and methodologies for describing social structure (e.g., Faust, 2006). These challenges and potential biases are compounded when focusing on group-level social structure beyond dyadic relationships, as describing such higher-order structures requires an additional layer of abstraction from the observable behaviour. If not interpreted critically, and with some understanding of their underlying processes, quantitative descriptions of animal social structures may therefore be biased by unintended or unknown confounds.

Significant advances in our understanding of social evolution and animal behaviour have been made in part due to Social Network Analysis (SNA), which provides a powerful tool to characterize and compare social phenotypes at different social scales, from individual to dyad to group (Farine & Whitehead, 2015). In recent years, SNA has largely become the default method for characterizing and describing animal social structure as it enables researchers to extract metrics for group-level structural properties, such as the density or modularity of the network (Farine & Whitehead, 2015; Sosa et al., 2021). However, while animal SNA continues to develop, with regular discussions and debates over best practices (Farine, 2024; Hart et al., 2022; Redhead et al., 2025), this approach does not come without limitations. In particular, SNA readily produces numeric estimates for group-level structural metrics from any behavioural data that researchers choose to represent as a network, without guaranteeing that these estimates are meaningfully comparable or representative of the underlying phenomena of interest (Sosa et al., 2021). Where the underlying processes underpinning emergent network structures are unknown, comparing global network metrics across different systems becomes particularly challenging as the reasons behind any detected differences remain opaque.

While the challenge of inferring true underlying patterns from limited observational data is well-known and targeted by several papers and packages on social network analysis, how these play out in shaping inference in group-based comparisons using social network methods has received less empirical attention. For example, the influence of group size on network metrics is inherent to the method: because many standard social network measures are defined in terms of the number of individuals and possible ties, changing group size inevitably shifts their expected values, even if the underlying social relationships remain identical (Anderson et al., 1999; Faust, 2006; Friedkin, 1981). Nonetheless, group size is rarely explicitly controlled or accounted for in comparative animal social network studies (Badihi et al., 2022; Balasubramaniam et al., 2018; Brooks et al., 2024; Parra et al., 2011; Pasquaretta et al., 2014; Rubenstein et al., 2007; Stroeymeyt et al., 2018), despite its potentially substantial impact on our interpretations and our ability to make meaningful comparisons.

Understanding the impact of individual, social, and ecological forces that underpin social interactions on their emergent social structures–that is the “content, quality, and patterning of social relationships emerging from repeated interactions between pairs [or larger sets] of individuals” (Kappeler, 2019)–along with the effects of observation biases, is crucial for accurate interpretation of any SNA. We refer to these three components (individual behaviour, external factors/ecology, and group structures) as the building blocks of social structures (Figure 1). If not interpreted critically, and with some understanding of their underlying processes, quantitative descriptions of animal social structures, like outputs from any quantitative methodology, may therefore be biased by unintended or unknown confounds. Indeed, group size and related demographic factors affect both social processes, data density, and the metrics we calculate, but we still lack a systematic understanding of the magnitude and form of these confounds.

**Figure 1.**
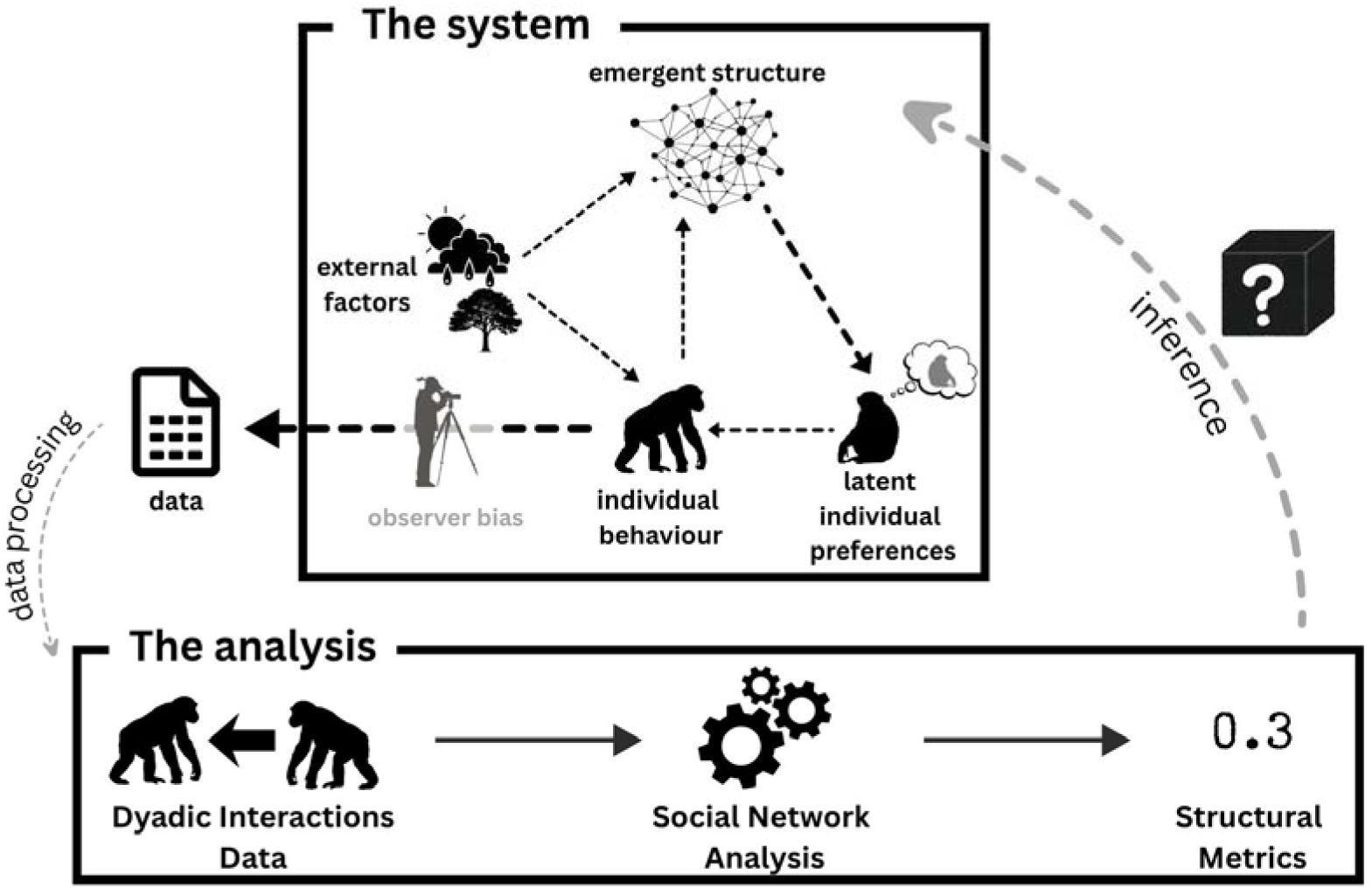
Conceptual model for inference on the evolution of social systems through social network analysis of observational data.

Comparative studies within and across species are becoming increasingly central to social evolution research, making it ever more important that we can provide reliable interpretations of social phenomena independently of stochastic demographic variation. Simulation-based approaches that estimate the predicted effects of such confounds on comparisons of social structure across groups offer a particularly promising way forward (Mielke et al., 2026; Palacios-Romo et al., 2019; Sueur, King, et al., 2011). They allow us to look beyond surface-level outputs of network metrics, but to consider how they compare to expected values driven by demographic and behavioural variation.

Our main objective is therefore to build a scalable model to simulate fission-fusion data, evaluate how SNAs are affected by potential confounds, and assess how this in turn shapes their ability to recover underlying social dynamics. We simulate fission-fusion spatial association networks to estimate the influence of multiple underlying confounds and sampling biases on commonly used social network metrics. Our approach builds a null model for the data collection process inspired by field observations of primates, specifically bonobos (*Pan paniscus*) and chimpanzees (*Pan troglodytes*).

## 2. Materials and Methods

### 2.1 Model

We simulated a basic model of social spatial association, inspired by *Pan* (chimpanzee and bonobo) fission-fusion dynamics (Kano, 1982; Nishida, 1968; Samuni et al., 2022; Surbeck et al., 2017), and designed to closely match with core principles of how empirical data are collected in real-time (ESM 1). Our simulations modelled groups of individuals, each with predefined social preferences (see 2.1.1) for every other group member, which determined how individuals tended to associate with one another and form temporary associations (i.e., parties). We simulated association data collected via scan sampling during daily focal follows, that is, following one “focal” individual across an entire day, during which probabilistic changes to that individual’s party composition were recorded over time. Each simulation run required three inputs: 1) the underlying dyadic (interindividual) association preferences (kept constant across the course of a given simulation run), 2) the target party sizes throughout the sampling period, 3) and focal individual for each day. We manipulated model parameters to simulate the socio-ecological constraints that act upon natural *Pan* systems (e.g., mean party size, party size variability). In doing so, we directly addressed questions about what social structures emerge under specific constraints, even when association was stochastic and non-deterministic.

#### 2.1.1 Model Inputs

1. **Interindividual preferences:** We represented dyadic association preferences with a symmetric matrix, meaning that in every dyad both individuals had equal attraction/repulsion towards the other. We initialized the value of dyadic preferences through different specified normal distributions, with either one or two peaks of varying spreads and differences between peaks (ESM 2.1, Figure S1), that can capture different interactional processes within social groups.
2. **Target party size:** We represented target party size with a matrix that stored the size of each observed party. In this matrix, each column corresponded to a day, each row to a scan (i.e., one observation of party composition), and each cell contained the size of the party for that scan. For the analysis reported here, we used a fixed setup of 24 scans per day, corresponding to a 12-hour focal follow with party composition recorded every 30 minutes (ESM 2.2).
3. **Focal identity:** Each day of each simulation run required one focal individual who was followed through party recombination events. As the observer follows a focal individual over the observation day, the focal individual appears in all parties recorded on that day, with the rest of the party determined by the combined preferences of all group members (see 2.1.2). Focal individuals were selected according to a specified sampling strategy (e.g., randomized, maximally equal, or biased by individual differences; see 2.3.2; ESM 2.7.2).

#### 2.1.2 Model runs

At the start of each simulated day, we created an initial party composition by sampling individuals based only on their dyadic association preferences for the focal individual. We extracted the focal individual’s preferences from the dyadic association matrix as a single vector and applied a softmax transformation, which takes a vector of real numbers and converts them to probabilities (all bounded between 0 and 1 and summing to 1), yielding the probabilities that a focal will be seen in the same party with each potential partner. Based on these probabilities, we then sampled as many individuals as required by the party matrix (minus one for the focal slot); partners with probabilities closer to 1 were more likely to be sampled.

We generated subsequent party compositions in multiple sequential steps based on the dyadic preferences on individuals within the current party. First, we calculated the focal’s average preference for i) the current party members and ii) all group members outside the party. These two mean values were converted, via the softmax function, into a probability that the focal either “stays” in or “leaves” the current party. We then drew one random sample using the softmax probabilities as weights for the two options (“stay” vs “leave”) to determine whether the focal “stays” or “leaves” the party. If the focal “left”, the next party was again formed in the same way as the day’s first party (i.e., by sampling from all group members based on their preference for the focal).

If the focal “stayed”, we updated the party membership as follows: for each non-focal member, we compared their average preference for members of the current party with their average preference for all individuals outside it. As with the focal, these two means were transformed with a softmax function to provide a stay/leave probability, where values closer to 1 reflect a high probability of staying, values close to 0 reflect a high probability of leaving. After all individuals had “decided” whether to stay or leave, using the same one-draw sampling method described for the focal, we adjusted the party to match the value specified in the party size matrix. If too many individuals decided to stay, we randomly removed individuals until the party matched the target size. If too few stayed, we filled the remaining slots by calculating, for all individuals outside the party, their average relative preference for the current (remaining) party members compared to the rest of the group, transforming these values via softmax, and sampling additional members according to the resulting probabilities. This procedure was repeated for each (possible) party composition change across the day (see ESM 2.3).

Each simulation run produced a data frame where each row corresponded to a single party and contained: i) the list of party members, ii) the identity of the focal individual, iii) the day number, iv) the scan number within that day, and v) the simulation run identifier (see example in Github_blinded).

### 2.2 Experimental design

We conducted four separate experiments, each utilizing an independent set of simulations with different parameters, to addressed different ways in which underlying features of individual attributes, dyadic interactions, and data collection biases may impact social network analysis for describing social structure of fission-fusion species. For all experiments, we ran 100 simulations per unique parameter combination (see ESM 2.4, Table S1), with a fixed rate of 24 party association scans per day, and an autocorrelation of 0.7 for the sizes of successive parties. All experiments swept group sizes from 5 to 155 in increments of 10. Each set of simulations followed the same sequence: 1) define the parameter space, 2) generate list of daily focal individuals, dyadic preferences, and party size matrices, 3) simulate the data collection procedure, 4) estimate edge weights for each dyad (simple ratio index [SRI; (Whitehead, 1995) ESM 2.4.1], and, in experiment 2, mean of posterior network [estimated via STRAND; Ross et al., 2024]; see ESM 2.4.2), 5) construct social networks using these estimated edge weights, and 5) calculate social network metrics. From networks built from each simulation output (i.e., one network per simulation run), we calculated five social network metrics (ESM 2.5): Density (mean edge weight across dyads), Clustering coefficient (global clustering using Barrat’s method; Barrat et al., 2004), Modularity (Lovain community detection Blondel et al., 2008), Edge CV (coefficient of variation of *edge weights*), and Node CV (coefficient of variation of *node strengths*). We compared metrics from simulated networks to networks obtained from published empirical datasets (Samuni et al., 2022; Surbeck et al., 2017; ESM 2.6)

### 2.3 Experimental conditions

#### 2.3.1 Experiment 1: Baseline Parameter Sweep

We first tested how group sizes and party sizes jointly influence measured social network metrics across a wide range of initialized dyadic preference distributions (ESM 2.7.1). For this experiment, all individuals were selected as focal for the same number of days (as much as possible).

#### 2.3.2 Experiment 2: Effect of individual-level gregariousness

We next examined how individual-level variation in behaviour shapes group-level network structure. We assessed how strongly our results depend on the assumptions that dyadic preference measures are independently sampled and that all group members are equally likely to be selected as focal (ESM 2.7.2). Because we target individual-level variation, for this experiment we also compared network metrics generated by using SRI as edge weight, to the metrics generated using the posterior network estimated via STRAND (a Bayesian model meant to account for sampling biases; Ross et al., 2024; ESM 5).

#### 2.3.3 Experiment 3: Clique structure

Next, we asked how social network analysis captures underlying clique (also referred to as subgrouping or clustering) structure in association preferences. By cliques we refer to stable association preferences between subsets of individuals from the wider group (ESM 2.7.3). We investigated how predefined clique structure in dyadic preferences, specifically the presence of tightly clustered cliques, influences SNA network metrics.

#### 2.3.4 Experiment 4: Observation time

Group size does not only influence social network analysis metrics, but also our ability to collect dense data on all individuals within the same time period. We therefore investigate how both absolute and relative observation time influence the consistency of network measures (ESM 2.7.4). We examined how the total number of observation days, both in absolute terms and relative to group size, affects measured social network metrics. In other words, given fixed dyadic preferences, we asked to what extent estimated network metrics converge as observation time increases.

## 3. Results

Through our simulated approach we directly tested how variation in specific features underlying fission-fusion systems impact SNA outputs. Across the four experiments, we generated 2 638 400 social networks (i.e., simulation runs) over 26 384 unique parameter combinations. Input dyadic preferences were correlated with SRI measured from simulated outputs, indicating that the simulated association data reasonably detected the pre-defined underlying dyadic association preferences (ESM 3).

### 3.1 Experiment 1. the impact of group size on social network metrics

All calculated SNA metrics showed non-linear relationships with group size, despite otherwise identical input parameters (n runs = 2 534 400). Runs with low overall variation in the sizes of parties (SD = 1) overlapped more closely with empirical estimates as compared to runs with high variation (SD = 7) in sizes of party and this pattern held across the full range of average party sizes and dyadic preference parameters (Figure 1). As expected, density showed a consistent pattern in relation to group size and tracked the density calculated from empirical data of similar group sizes. Other metrics also changed systematically with group size, but exhibited greater variation within each group size (i.e., wider confidence intervals). All runs substantially underestimated the node coefficient of variance (i.e., the degree of heterogeneity of individual total connections) relative to the empirical data, most notably in larger groups (Figure 2).

**Figure 2.**
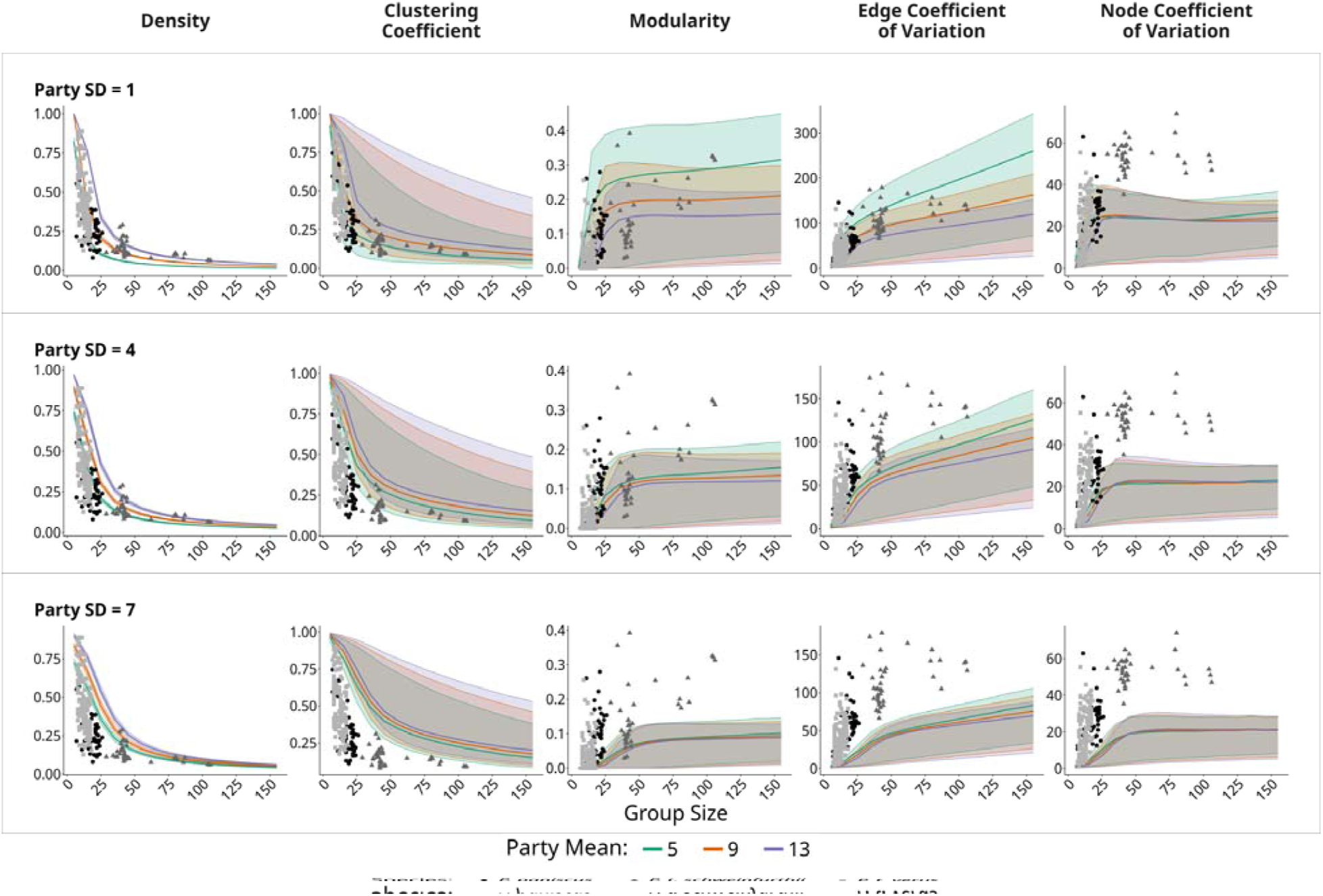
Sensitivity of social network metrics across different group and party sizes. Changes in social network metrics calculated across simulated runs with varying mean party size (Party Mean), party size variability (Party Standard Deviation: top = low variability, bottom = high variability) and groups of different sizes (n runs = 2 534 400; ESM 2.7.1). See ESM 4 (Figure S2) the effects of group and party sizes on each SNA metric relative to density. Points represent SNA metrics calculated from published empirical data (Samuni et al., 2022; Surbeck et al., 2017; ESM 6.1; Tables S2-4 for quantitative comparisons).

### 3.2 Experiment 2. the impact of individual-level variation on social network metrics

Introducing individual-level variability in “gregariousness” (i.e., baseline attraction towards other individuals to which dyadic preferences are added) and focal selection strategy (i.e., assuming individuals are not all selected as focal equally often) jointly influenced social network metrics (n runs = 25 600). When individuals varied, even to a small degree, in gregariousness (coefficient of variance = 0.1) we observed a moderate impact on the SNA metrics across group sizes (Figure 3), particularly in edge and node CV. This influence was amplified when we increased heterogeneity of gregariousness between individuals (coefficient of variance = 0.4). Without including the gregariousness condition, our simulations consistently resulted in lower node CV as compared to empirical networks of similar size. When we introduced the gregariousness condition, node CV in simulated networks increased, and in conditions with high heterogeneity in gregariousness between individuals (greg_cv = 0.4), simulated networks indicated higher node CV than empirical networks of similar size.

**Figure 3.**
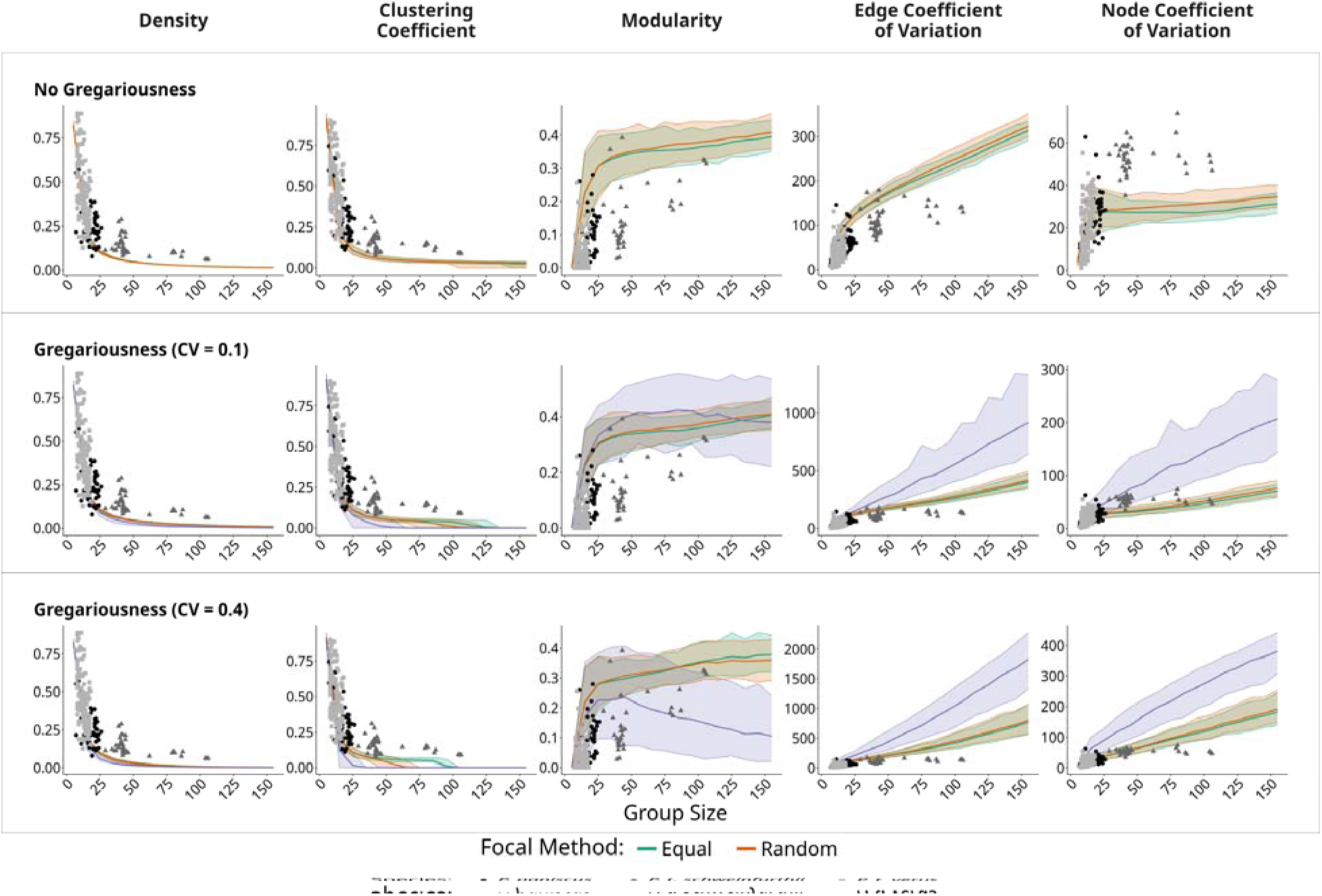
Sensitivity of social network metrics across different group and party sizes with changing individual-level gregariousness and focal sampling method. Changes in social network metrics calculated across simulation runs with varying degree of heterogeneity in individual-level gregariousness (top = no variation in gregariousness; middle = low heterogeneity, coefficient of variance = 0.1; top = high heterogeneity, coefficient of variance =0.4), across different focal sampling methods (equal; random; weighted by individual gregariousness), and groups of different sizes (n runs = 25 60; ESM 2.7.2). Points represent SNA metrics calculated from published empirical data (Samuni et al., 2022; Surbeck et al., 2017; ESM 6.2; Tables S5–7 for quantitative comparisons).

Using a fully randomized sampling strategy (i.e., selecting each focal independently instead of systematically ensuring equal coverage to all individuals) had minimal effects on the SNA metrics analyzed as compared to the equal sampling method. In contrast, when more gregarious individuals were more likely to be chosen as focal, this bias substantially altered all SNA metrics except network density. Gregariousness-weighted focal selection produced substantially higher node- and edge-level CV. Modularity did not change uniformly across group sizes under the gregariousness-weighted focal selection condition; it exhibited a non-linear response first increasing with increase in group sizes up to around 50 individuals and then decreasing and was characterized by greater variability (i.e., wider confidence intervals).

### 3.3 Experiment 3. the impact of clique structure on social network metrics

When imposed a clique-based social structure to the dyadic preferences underlying the simulation, we found that the SNA metrics were influenced by an interaction between group size and the number of cliques introduced (n runs = 27 200; Figure 3). While density remained relatively constant across different numbers of cliques (except for the smallest groups where many observed parties include all group members), clustering coefficient, modularity, and both edge- and node-level CV were altered by clique structure (see columns in heatmaps of Figure 4). We observe that a uniform dyadic preference matrix, where all dyads were equally attracted to one another, produced relatively low modularity, edge- and node-level CV, compared to dyadic preference matrices with clique structures. Furthermore, across group sizes, the clustering coefficient was high in the uniform dyadic preference condition relative to both previous simulation experiments and relative to the empirical data.

**Figure 4.**
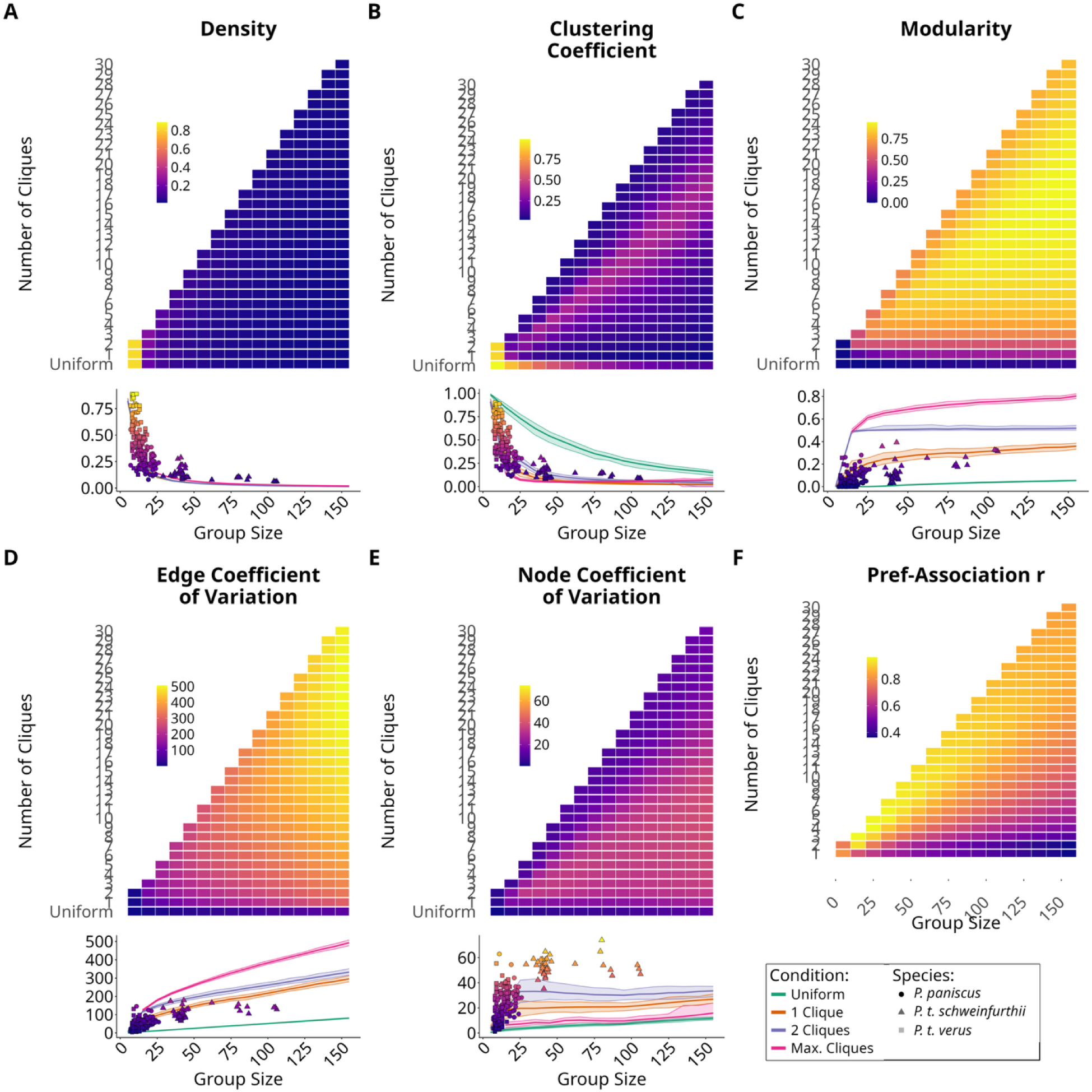
Clique structures exert non-linear effects on network metrics. Impact of initialized subgroup structure preferences on social network metrics across groups of different sizes (n runs = 27 200; ESM 2.7.3). Uniform condition refers to a fully uniform initial dyadic preferences (i.e., a matrix of all zeros) while clique conditions refer to dyadic preferences sampled from a distinct normal distribution for within-cloque and between-clique dyads (i.e., 1-clique condition all dyads sampled from the same normal distribution). Points represent SNA metrics calculated from published empirical data (Samuni et al., 2022; Surbeck et al., 2017; ESM 6.3; Tables S8–9 for quantitative comparisons); point colour use the same scale as the simulated data heatmaps.

We observe peaks in several SNA metric measures for some combinations of group size and number of cliques (e.g., diagonal band in clustering coefficient, Figure 4B), with values decreasing as conditions move away from these optima. For example, clustering coefficient was highest along a band where clique size was approximately 10 individuals, with lower values for both smaller and larger cliques. Modularity was lower for very large or very small cliques than for intermediate-sized ones. Simulations based on non-uniform dyadic preferences produced social network metrics that more closely matched the empirical estimates than did simulations with uniform dyadic preference. However, within non-uniform conditions, scenarios with multiple distinguishable cliques were further from empirical estimates than those without. Across all clique structures, node CV remained consistently lower in the simulated model outputs as compared to the empirical data; however, in this case, the two-clique condition most closely tracked the empirical data.

### 3.4 Experiment 4. the impact of observation protocols on SNA metrics

Observation periods with fewer total observation days and fewer observation days per focal had little impact on density and clustering coefficient estimates, though showed greater variability than observation periods with more observation days. In contrast, lower observation effort (both absolute and per focal) tended to produce higher measures of network modularity, edge CV, and node CV than the same population observed for more days (n runs = 51 200). This effect was most prominent in 30-day observation windows. Further, higher modularity, edge CV, and node CV measures observed in the 30-day observation condition (as compared to 90-, 180-, and 360-day) became increasingly prominent with increasing group size. Still, SNA metric values began to converge within 90-180 observation days (with unbiased focal sampling) – even at the largest group sizes, where not all individuals are selected as focal. Network modularity calculated from one-year periods of empirical data were typically closer than were 3-month observation periods to the simulated curves, although there were relatively few one-year empirical observation periods. As in our other experiments, node CV was consistently lower as compared to empirical data, regardless of observation period duration (discussed in experiment 1). However, we observe higher node- and edge-level CV in simulation runs with fewer observation days (e.g., 30 days or 0.5 days per individual), suggesting that data sparsity can inflate measures of variance (Figure 5).

**Figure 5.**
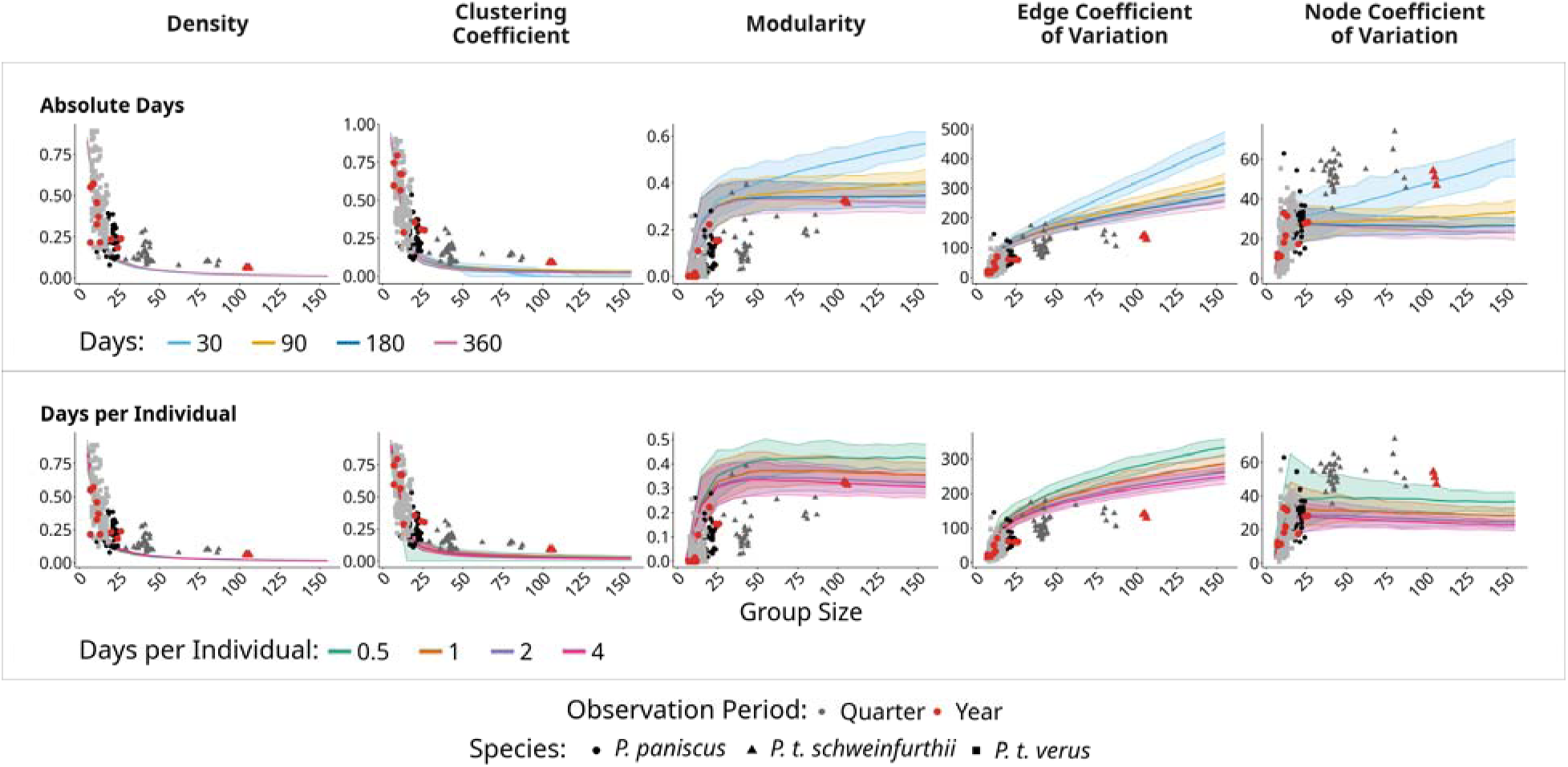
Observation effort influences social network metrics. Impact of absolute observation time (top row, number of days = 30, 90, 180, 360) and number of observation days relative to group size (bottom row, n = 0.5, 1, 2, 4) on measured social network metrics across groups of different sizes (n runs = 51 200, ESM 2.7.4). Points represent SNA metrics calculated from published empirical data (Samuni et al., 2022; Surbeck et al., 2017; ESM 6.4; Tables S10–11 for quantitative comparisons).

## 4. Discussion

Our model demonstrates how multiple levels of confounds can influence outputs from SNA, highlighting the challenges with inferring group-level structural properties and pressures from observational social network data. We show that demographic and association constraints, latent individual social dispositions, and observation bias all have non-linear but consistent effects on SNA metrics commonly used to describe group-level properties both when using SRI and STRAND methods. This model highlights the wider potential of simulation approaches for addressing and quantifying the relationships between latent and emergent features of complex systems, taking a critical step in understanding the evolutionary mechanisms of group-level social structures.

### External factors

Group size consistently had a prominent effect on measurements derived from SNA. While differences in group size may themselves reflect meaningful socio-ecological variation, we show that they may also confound attempts to compare the structural properties of differently sized groups. The strongest effects were observed for density, which consistently decreased with group size regardless of variation in other model parameters. Because density is closely related to the ratio of average party size to total group size, this relationship is not surprising (Faust, 2006), but further illustrates that density should not be used to characterize anything beyond that simple demographic network property. Nevertheless, the effects of group size on other SNA metrics remained even when accounting for differences in density (ESM 4). Effects of group size across SNA metrics were especially prominent in smaller groups, where even modest changes in group size can substantially alter calculated network measures. These effects were compounded in “raw” social network metric values without accounting for density. The strong effects of group size across SNA metrics, particularly at the lower end of the group size range, indicates that common demographic changes, such as births, deaths, or migrations, as well as analytical decisions about network inclusion criteria (e.g., age; observation effort; Fedurek & Lehmann, 2017), can produce dramatically different metric values. Consequently, temporal variation in network metrics may reflect changes in group size or composition, rather than changes in social behavior and decision-making alone.

Group size is not only a confound but is also expected to have real impacts on behavioral patterns that researchers seek to measure. For example, coordinating travel and foraging decisions is likely more challenging in larger groups. Individuals may respond by selectively prioritizing a subset of group members to travel or forage with (e.g., those with whom they share stronger social relationships). Indeed, across simulated and empirical networks we found that larger groups exhibited higher modularity, more variation in strength of dyadic associations, and more variation in individuals’ centrality within the network (also relative to density, ESM 4). Yet, even within the same group size and party sizes, there was variation around the means of network metrics in both simulated and empirical data. This variance may be related to real structural differences between groups detected by these SNA measures (e.g., due to different distributions of dyadic preferences), but is uninterpretable in isolation without careful consideration of the confounding effects of group sizes and party sizes on these measures.

While group size represents one important axis of the social environment that interacts with social structure, societies also experience flexible within-group association patterns that add another layer of demographic fluctuations. Even when ecological constraints limit party or unit size, groups can be large as individuals can maintain contact with all group members through reassorting into parties of varying composition (high-level fission-fusion) or by organizing into stable, nested social units (multi-level). In societies characterized by high-level fission fusion, like chimpanzees and bonobos, evolutionary pressures on overall group size are thus at least partially distinct from the evolutionary pressures on foraging party size, as long as the group can maintain a sufficient level of connectivity between group members (Sandel et al., 2026). The average party sizes chosen for our simulations (5, 9, 13) are within the range of reported average party sizes of wild chimpanzees and bonobos (Kano, 1982; Samuni et al., 2022), and are therefore representative of real-world social dynamics. Using these initial party size parameters, we found that groups with smaller parties relative to their overall group size had higher modularity and more heterogeneous dyadic association strengths (also relative to density, ESM 4), meaning that individuals were less connected within these groups. When average party size was held constant, groups parameterized to have greater variability in party sizes (i.e., party_sd) were more broadly connected (less modular) and contained more homogenous inter-dyadic ties (more equally distributed relationships) than groups parameterized to have more consistent party sizes (also relative to density, ESM 4). Our simulations suggest that distinguishing the independent and interacting effects of average party size and variability is essential for understanding variation in group-level social structure across species and populations.

### Individual behaviour

Social structure emerges from individual behaviour and interactions among group members, with individuals differing in their ability and propensity to interact with others. Using simulations we could explore assumptions related to such individual differences, thereby recognising the contribution of inter-individual differences in behaviour for emerging social structures (De Moor, Hart, et al., 2025; Hinde, 1976; Kawam et al., 2025; Neumann & Fischer, 2026). One way we achieved this was by varying the attraction of individuals to others (i.e., ‘gregariousness’) and therefore their likelihood to be recorded in parties. We used ‘gregariousness’ as a proxy with which to test whether greater heterogeneity in individual behaviour also leads to more heterogeneous group structures detected through SNA. As expected, when we simulated greater inter-individual variability in gregariousness, social networks exhibited greater heterogeneity in the strength of both individual (higher node CV) and dyadic (higher edge CV) connectedness. Nevertheless, even without simulating inter-individual differences in gregariousness, and with group and party size model parameters that mimic real-world data, larger groups (i.e., greater partner availability) were associated with more heterogeneous individual and dyadic connections, reinforcing the idea of group size as a potential component of social differentiation and complexity (Steiner, 1966; Sueur, Deneubourg, et al., 2011). In this sense, simulations become particularly useful for understanding and exploring assumptions about how latent sources of variation, here gregariousness, may impact measures of social structure. Many additional differences exist between individuals of the same social group, for example in sex, age, kinship, reproductive state, and dominance relationships; future implementations of our model could specify such individual attributes and their expected consequences, and test their effects on social structure. Such simulation work may help inform evolutionary theory about how selective pressures on social structure can act through individual behaviour and their latent preferences.

### Subgrouping and social connectivity

Larger groups increase the number of social relationships that individuals must manage, potentially increasing uncertainty, and with that, the cognitive demands of sociality (e.g., Byrne & Whiten, 1988). Stable subgrouping may reduce these demands by making relationships more predictable (Ramos-Fernandez et al., 2018). For example, in some high-level fission-fusion systems, large group size has been associated with the emergence of subgrouping structures (Badihi et al., 2022; Mitani & Amsler, 2003). We therefore represented potential subgrouping by imposing higher-order association preferences, such that dyadic preferences were not independent but were structured into ‘cliques’ which represent stable subgroups in which individuals preferentially associate within the larger group. This enables us to probe the extent of how high-level fission-fusion association patterns could be expected to change in settings with complete partitioning of preferences.

Our simulations suggest a non-linear and non-monotonic relationship between the number of cliques imposed and network structure, where individual and dyadic association strengths in groups with either few or many cliques were more homogeneous. This pattern arises from the relation between clique size and average party size: when parties are generally smaller than cliques, individuals are less likely to interact with *all* other clique members; when parties are generally larger than cliques, they are more likely to interact with non-clique members. Higher modularity may reflect greater predictability in spatial association, as individuals are more likely to form parties composed (only) of individuals from the same clique. Taken to the extreme, stable partitions of a larger group may begin to resemble the strategies used in multi-level societies. Depending on socio-ecology, increased predictability through stable subgrouping may come at the cost of cohesion across the wider group, potentially increasing the risk of permanent (and potentially lethal) fission along clique boundaries (e.g., Sandel et al., 2026). The modularity values observed from empirical *Pan* datasets most closely resembled that of simulations in which no stable cliques were imposed but individuals/dyads differed in their social preference. This was also true for the largest community in the empirical dataset. Therefore, our findings suggest that real *Pan* groups maintain differentiated social relationships without forming fully discrete, stable cliques. In other words, they may tolerate social uncertainty to preserve cohesion across the group and reduce the risk of permanent fission.

### Observation bias and effort

Our inference of animal behaviour is ultimately processed through the lens of our observation biases and effort (reviewed in Webster & Rutz, 2020). Several works have targeted how observed behavioural data relates to the underlying interactions and latent preferences of individuals, but have largely focused on dyadic relationships (Mielke et al., 2026, but see Davis et al., 2018). Challenges to inference due to incomplete or biased data are compounded when addressing questions of group-level social structure due to the nonlinear scaling of dyadic (and higher-order) relationships as group size increases. This is uniquely salient for association data which is inherently higher-order (e.g., an individual C may associate with A as a byproduct of seeking to associate with B, if A and B preferentially associate). Even when a single individual is followed as a focal, association data captures more than that individual’s decisions, such that all other associations observed in the party are recorded simultaneously.

Our simulations show that even for data that is inherently higher-order and allows for the simultaneous monitoring of multiple individuals at a time (like association data), observation effort influences SNA metrics and therefore sampling windows must be chosen carefully. In our experiment, SNA metrics stabilized (i.e., additional data provided diminishing returns) at an observation duration of at least 180 days, or around 2 days per individual under a relative sampling regime. Adding more observation days after this threshold did not change the metrics calculated using SNA. These patterns provide further proof that our ability to reliably capture global social structures is contingent on observation effort, a dependence that is likely to be amplified when the behavior of interest is predominantly dyadic (e.g., grooming). Even with high effort, observational biases can confound inference, for example, when ‘gregarious’ individuals are followed more often (Experiment 2). Our simulated model provides a tool with which to estimate the minimum sampling time to reach replicable estimates of social structure, given a set of assumptions about the system and observational constraints.

Nevertheless, our model did not account for the fact that association patterns can fluctuate temporally within a given system. Seasonal variation can lead to fluctuations in how individuals interact (e.g., due to variation in food availability and distribution), leading to consistent changes in mean and variability of party sizes. Researchers should consider not just how much, but also when they are sampling. Real seasonal variation in *Pan* systems may lead to differences in party sizes (and variability in party sizes) throughout the year according to seasonal resource availability. Although we do not simulate temporal variation over distinct and changing seasons, larger party size variability can be understood to capture some element of seasonal fluctuations in party sizes over a given time period. Greater variability in party sizes (i.e., more than our simulated SD = 1), likely requires longer observation effort for SNA metrics to stabilize, even when individuals have identical latent preferences. Conversely, larger average parties relative to group size may require shorter observation effort to stabilize. When choosing appropriate observation windows, researchers should consider such levels of variation as they likely impact the time table to reach the most accurate descriptions of social structure for a given time period, and future simulation work may be able to generate more precise recommendations.

## Conclusion

Across the board, group size was a major driver of changes observed in social network metrics. This pattern was strongest at small group sizes, as expected, given that social network metrics often reflect a relationship between group size and interaction opportunities. More specifically, in flexible fission-fusion systems the ratio of party size to group size reflects the ability of groupmates to encounter all other individuals and therefore the proportion of dyadic connections that can be realized simultaneously. With constant party size, the addition of even one individual to a small group has a more dramatic effect on this ratio. However, even when we controlled for density (i.e., the relationship between party size and group size), group size still had a pronounced effect on SNA metrics suggesting that group size can confound our measures of social structure beyond what can be explained by changes in density. When designing studies to investigate the drivers of group-level structures, researchers must make careful decisions about how to account for or integrate group-size into the research question.

Can group size be considered as a driver of specific group-level structures or is group size a mechanism by which groups adapt to achieve an optimal structure that was selected for by other evolutionary or proximate pressures? We show that different, and sometimes opposite, underlying processes can lead to similar outputs from SNA, making it difficult to determine the drivers of particular social structures through SNA metrics. With growing availability of large comparative datasets in animal behaviour studies and access to statistical tools such as SNA (De Moor, Skelton, et al., 2025; Sah et al., 2024), researchers should strive to fully understand the limitations and underlying mechanisms behind the tools they apply. This is especially important for tools, such as social network analysis, that enable the quantification and description of higher order structures that are extracted from underlying individual behaviour and interactional processes. While we focused on how changes to a fission-fusion system can influence SNA, other methodologies and analysis techniques can be applied to the same simulation outputs to test their sensitivity to similar (or different) changes in a study system. The results of our simulations are full party composition scans, and so any measures relating to party composition (e.g., hyper-graphs, entropy), can be targeted with such simulated data.We advocate for systematic simulation studies as necessary methodological advances before attempting to infer group-level selective pressures on animal social systems. Overall, our model provides a simple and scalable tool to analyze how global network properties are affected by low-level preferences, constraints, and biases in a fission-fusion system.

## Author Contributions

All authors conceived the ideas and designed methodology; James Brooks and Gal Badihi analysed the data; all authors led the writing of the manuscript; all authors contributed critically to the drafts and gave final approval for publication.

## Supporting information

ESM

## Acknowledgements

We thank Prof. Daniel Redhead for providing valuable comments on our methods and results, and the Cooperative Cultures Research Group for feedback and discussion. We thank the Diverse Intelligences Summer Institute for providing the setting where these ideas were first discussed and for an Early Career Collaboration Enhancement grant that enabled us to conduct this study. Finally, we thank the authors of Surbeck et al. (2013) and Samuni et al. (2022) for making their data open access. We used Large Language Models (Claude and ChatGPT) to aid code and repository formatting; all original code used in our model simulation was written and verified by the authors prior to AI-assisted formatting.

## Funding

Diverse Intelligence Summer Institute, Early Careers Collaboration Enhancement grant, Emmy Noether Program (513871869), Deutsche Forschungsgemeinschaft (DFG), Alexander van Humbold Foundation, Max Planck Society

## Data availability statement

Empirical data were obtained from open access sources (Surbek et al., 2017; Samuni et al., 2022). Simulated datasets and model scripts can be sourced at https://github.com/Cooperative-Evolution/Simulating_social_structures and all data presented in results can be found at doi.org/10.5281/zenodo.22746597

## Conflicts of Interest

We declare no conflicts of interest.

## Notes

### Competing Interest Statement

The authors have declared no competing interest.

### Summary of Updates

formatting was revised to meet the requirements of submission

## References

1. Anderson, B. S., Butts, C., & Carley, K. (1999). The interaction of size and density with graph-level indices. Social Networks, 21(3), 239–267. 10.1016/S0378-8733(99)00011-8

2. Badihi, G., Bodden, K., Zuberbühler, K., Samuni, L., & Hobaiter, C. (2022). Flexibility in the social structure of male chimpanzees (Pan troglodytes schweinfurthii) in the Budongo Forest, Uganda. R. Soc. Open Sci., 9, 220904. 10.1098/rsos.220904

3. Balasubramaniam, K. N., Beisner, B. A., Berman, C. M., De Marco, A., Duboscq, J., Koirala, S., Majolo, B., MacIntosh, A. J., McFarland, R., Molesti, S., Ogawa, H., Petit, O., Schino, G., Sosa, S., Sueur, C., Thierry, B., De Waal, F. B. M., & McCowan, B. (2018). The influence of phylogeny, social style, and sociodemographic factors on macaque social network structure. American Journal of Primatology, 80(1), e22727. 10.1002/ajp.22727

4. Barrat, A., Barthélemy, M., Pastor-Satorras, R., & Vespignani, A. (2004). The architecture of complex weighted networks. Proceedings of the National Academy of Sciences, 101(11), 3747–3752. 10.1073/pnas.0400087101

5. Brooks, J., Maeda, T., Ringhofer, M., & Yamamoto, S. (2024). Oxytocin homogenizes horse group organization. iScience, 27(7), 110356. 10.1016/j.isci.2024.110356

6. Byrne, R. W., & Whiten, A. (1988). Machiavellian intelligence: Social expertise and the evolution of intellect in monkeys, apes, and humans. Clarendon Press.

7. Davis, G. H., Crofoot, M. C., & Farine, D. R. (2018). Estimating the robustness and uncertainty of animal social networks using different observational methods. Animal Behaviour, 141, 29–44. 10.1016/j.anbehav.2018.04.012

8. De Moor, D., Hart, J. D. A., Franks, D. W., Brent, L. J. N., Silk, M. J., & Brask, J. B. (2025). Latent layers in social networks and their implications for comparative analyses. Behavioral Ecology, 36(6), araf113.

9. De Moor, D., Skelton, M., MacaqueNet, Amici F., Arlet, M. E., Balasubramaniam, K. N., Ballesta, S., Berghänel, A., Berman, C. M., Bernstein, S. K., Bhattacharjee, D., Bliss Moreau, E., Brotcorne, F., Butovskaya, M., Campbell, L. A. D., Carosi, M., Chatterjee, M., Cooper, M. A., Cowl, V. B., … Brent, L. J. N. (2025). MacaqueNet: Advancing comparative behavioural research through large scale collaboration. Journal of Animal Ecology, 94(4), 519–534. 10.1111/1365-2656.14223

11. Farine, D. R. (2024). Modelling animal social networks: New solutions and future directions. Journal of Animal Ecology, 93(3), 250–253. 10.1111/1365-2656.14049

12. Farine, D. R., & Whitehead, H. (2015). Constructing, conducting and interpreting animal social network analysis. Journal of Animal Ecology, 84(5), 1144–1163. 10.1111/1365-2656.12418

13. Faust, K. (2006). Comparing social networks: Size, density, and local structure. Advances in Methodology and Statistics, 3(2), 85–216. 10.51936/sdbv3216

14. Fedurek, P., & Lehmann, J. (2017). The effect of excluding juveniles on apparent adult olive baboons (Papio anubis) social networks. PLOS ONE, 12(3), Article 3. 10.1371/journal.pone.0173146

15. Friedkin, N. E. (1981). The development of structure in random networks: An analysis of the effects of increasing network density on five measures of structure. Social Networks, 3(1), 41–52. 10.1016/0378-8733(81)90004-6

16. Hart, J. D. A., Weiss, M. N., Brent, L. J. N., & Franks, D. W. (2022). Common permutation methods in animal social network analysis do not control for non-independence. Behavioral Ecology and Sociobiology, 76(11), 151. 10.1007/s00265-022-03254-x

17. Hinde, R. A. (1976). The Use of Differences and Similarities in Comparative Psychopathology. In G. Serban & A. Kling (Eds), Animal Models in Human Psychobiology (pp. 187–202). Springer US. 10.1007/978-1-4684-2184-2_11

18. Kano, T. (1982). The social group of pygmy chimpanzees (Pan paniscus) of Wamba. Primates, 23(2), Article 2. 10.1007/BF02381159

19. Kappeler, P. M. (2019). A framework for studying social complexity. Behavioral Ecology and Sociobiology, 73(1), 13. 10.1007/s00265-018-2601-8

20. Kawam, B., Ostner, J., McElreath, R., Schülke, O., & Redhead, D. (2025). A causal framework for the drivers of animal social network structure. PLoS Comput Biol, 21(9), e1013370. doi.org/10.1371/journal.%20pcbi.1013370

21. Mielke, A., Testard, C., Motes-Rodrigo, A., Brent, L. J. N., & De Moor, D. (2026). Observation methods in animal behaviour: A simulation study of performance. Animal Behaviour, 237, 123604. 10.1016/j.anbehav.2026.123604

22. Mitani, J., & Amsler, S. (2003). Social and spatial aspects of male subgrouping in a community of wild chimpanzees. Behaviour, 140(7), 869–884. 10.1163/156853903770238355

23. Neumann, C., & Fischer, J. (2026). (De)composing sociality: Disentangling individual-specific from dyad-specific propensities to interact. Methods in Ecology and Evolution, 17, 963–978. 10.1111/2041-210x.70239

24. Nishida, T. (1968). The social group of wild chimpanzees in the Mahali Mountains. Primates, 9(3), 167–224. 10.1007/BF01730971

25. Palacios-Romo, T. M., Castellanos, F., & Ramos-Fernandez, G. (2019). Uncovering the decision rules behind collective foraging in spider monkeys. Animal Behaviour, 149, 121–133. 10.1016/j.anbehav.2019.01.011

26. Parra, G. J., Corkeron, P. J., & Arnold, P. (2011). Grouping and fission–fusion dynamics in Australian snubfin and Indo-Pacific humpback dolphins. Animal Behaviour, 82(6), 1423–1433. 10.1016/j.anbehav.2011.09.027

27. Pasquaretta, C., Levé, M., Claidière, N., van de Waal, E., Whiten, A., MacIntosh, A. J. J., Pelé, M., Bergstrom, M. L., Borgeaud, C., Brosnan, S. F., Crofoot, M. C., Fedigan, L. M., Fichtel, C., Hopper, L. M., Mareno, M. C., Petit, O., Schnoell, A. V., di Sorrentino, E. P., Thierry, B., … Sueur, C. (2014). Social networks in primates: Smart and tolerant species have more efficient networks. Scientific Reports, 4(1), 7600. 10.1038/srep07600

28. Ramos-Fernandez, G., King, A. J., Beehner, J. C., Bergman, T. J., Crofoot, M. C., Di Fiore, A., Lehmann, J., Schaffner, C. M., Snyder-Mackler, N., Zuberbühler, K., Aureli, F., & Boyer, D. (2018). Quantifying uncertainty due to fission–fusion dynamics as a component of social complexity. Proceedings of the Royal Society B: Biological Sciences, 285(1879), 20180532. 10.1098/rspb.2018.0532

29. Redhead, D., Kawam, B., Young, J.-G., Franks, D., Philson, C. S., Duijn, M. van, Hart, J., McElreath, M. B., McElreath, R., Power, E. A., Ross, C., Sosa, S., Steglich, C., Weiss, M., & Brent, L. J. N. (2025). Five misunderstandings in animal social network analysis. https://ecoevorxiv.org/repository/view/9817/

30. Ross, C. T., McElreath, R., & Redhead, D. (2024). Modelling animal network data in R using STRAND. Journal of Animal Ecology, 93(3), 254–266. 10.1111/1365-2656.14021

31. Rubenstein, D. I., Sundaresan, S., Fischhoff, I., & Saltz, D. (2007). Social Networks in Wild Asses: Comparing Patterns and Processes among Populations. Erforsch. Biol. Ress. Mongolei, 10, 159–176.

32. Sah, P., Collier, M., Ali, S., Mendez, J., & Bansal, S. (2024). *Animal Social Network Repository* (Version v2.01) [Data set]. 10.5281/zenodo.14009255

33. Samuni, L., Langergraber, K. E., & Surbeck, M. H. (2022). Characterization of Pan social systems reveals in-group/out-group distinction and out-group tolerance in bonobos. Proceedings of the National Academy of Sciences, 119(26), e2201122119. 10.1073/pnas.2201122119

34. Sandel, A. A., He, Y., Ren, J., Kei, Y. L., Lee, K. C., Clark, I. R., Reddy, R. B., Negrey, J. D., Birungi, C., Apamaku, B. A., Kanweri, D., Kalunga, D., Aliganyira, C., Ramírez-Amaya, S., Nakayima, P., Katumba, R., Kamugyisha, B., Acosta-Florez, D., van Boekholt, B., … Mitani, J. C. (2026). Lethal conflict after group fission in wild chimpanzees. Science, 392(6794), 216–220. 10.1126/science.adz4944

35. Siracusa, E. R., Bal, X., Moor, D. D., Albery, G. F., Beattie, R., Ravindran, S., Wiersma, E., Wilson, A. R., Pemberton, J. M., Nussey, D. H., & Silk, M. J. (2026). Environment-dependent benefits of sociality in Soay sheep. Proc. R. Soc. B, 293, 20252821.

36. Sosa, S., Sueur, C., & Puga Gonzalez, I. (2021). Network measures in animal social network analysis: Their strengths, limits, interpretations and uses. Methods in Ecology and Evolution, 12(1), 10–21. 10.1111/2041-210X.13366

37. Steiner, I. D. (1966). Models for Inferring Relationships Between Group Size and Potential Group Productivity. Behavioral Science, 11(4), 273–283.

38. Stroeymeyt, N., Grasse, A. V., Crespi, A., Mersch, D. P., Cremer, S., & Keller, L. (2018). Social network plasticity decreases disease transmission in a eusocial insect. Science, 362(6417), 941–945. 10.1126/science.aat4793

39. Sueur, C., Deneubourg, J.-L., Petit, O., & Couzin, I. D. (2011). Group size, grooming and fission in primates: A modeling approach based on group structure. Journal of Theoretical Biology, 273(1), 156–166. 10.1016/j.jtbi.2010.12.035

40. Sueur, C., King, A. J., Conradt, L., Kerth, G., Lusseau, D., Mettke-Hofmann, C., Schaffner, C. M., Williams, L., Zinner, D., & Aureli, F. (2011). Collective decision-making and fission-fusion dynamics: A conceptual framework. Oikos, 120(11), 1608–1617.

41. Sueur, C., Solé, R., & Deneubourg, J.-L. (2026). Collective social niche construction shaping adaptive social networks. Trends in Ecology & Evolution, 41(7), 607–617. 10.1016/j.tree.2026.04.001

42. Surbeck, M., Girard-Buttoz, C., Boesch, C., Crockford, C., Fruth, B., Hohmann, G., Langergraber, K. E., Zuberbühler, K., Wittig, R. M., & Mundry, R. (2017). Sex-specific association patterns in bonobos and chimpanzees reflect species differences in cooperation. Royal Society Open Science, 4(5), 161081. 10.1098/rsos.161081

43. Webster, M. M., & Rutz, C. (2020). How STRANGE are your study animals? Nature, 582(7812), 337–340. 10.1038/d41586-020-01751-5

44. Whitehead, H. (1995). Investigating structure and temporal scale in social organizations using identified individuals. Behavioral Ecology, 6(2), 199–208. 10.1093/beheco/6.2.199

