## Supplementary material for "The building blocks of social structure: simulating constraints of social network analysis for inference about group-level properties": ESM

**ESM 1. Target system**

Fission-fusion association dynamics are phylogenetically widespread in the animal kingdom, observed in species from primates to birds to cetaceans, and occur in varying degrees and in multiple forms (e.g., Madsen & De Silva, 2024). In systems with high-level fission-fusion, group members form, split, and reassort into dynamic subgroups of variable size, composition, and duration (Aureli et al., 2008). For example, in chimpanzees and bonobos, groups (or “communities”) are rarely all together and instead individuals move between and associate in subgroups (referred to as “parties”) of variable size and composition over the course of a day (Kano, 1982; Nishida, 1968; Samuni et al., 2022; Surbeck et al., 2017).  Due to its flexible and variable form, fission-fusion is central to understanding many social systems, but remains difficult to capture and compare in robust structural terms (Madsen & De Silva, 2024), partially due to confounding effects of observation effort, temporal scale, and demographic variation.

**ESM 2. Model Parameters and Runs**

***2.1 Distributions of individual preferences in dyadic preference matrices***

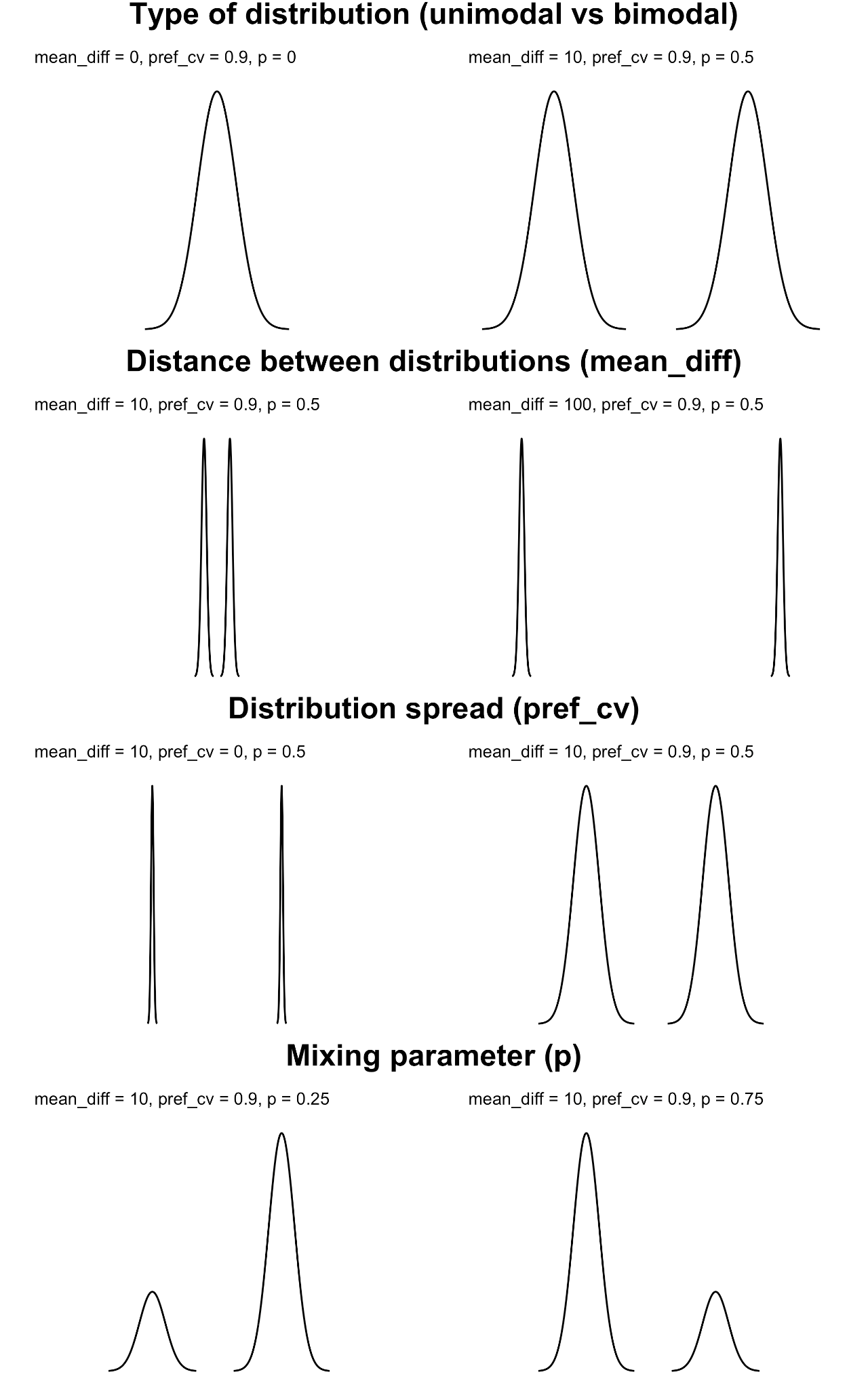

**Figure S1.** *Representation of preference distributions used to generate dyadic preferences matrix. While not an exhaustive list this figure illustrates the relative effects of changing distance between the two bimodal peaks (mean difference), the spread of the distributions (preference CV), and bias to sample from one distribution over the other (mixing parameter), used to change the shape of the distribution(s).*

***2.2. Party size matrix generation:***

The matrix was generated from three inputs: the average party size, the variability in party size, and the strength of autocorrelation between consecutive party scans. Given a chosen average party size, we set the time series of party sizes for each day using an ARIMA (Autoregressive Integrated Moving Average) process, which generates a time series centered around a specified mean with given standard deviation and autocorrelation between successive elements. The ARIMA process was set up to i) maintain a specified autocorrelation between the sizes of successive party scans (kept at 0.7 throughout runs presented here to provide a strong positive correlation between consecutive party sizes while allowing some fluctuation, as is typical of real fission-fusion systems [Kano, 1982; Nishida, 1968; Samuni et al., 2022]) and ii) produce a prescribed level of variability (standard deviation) around predefined average party sizes across all generated parties. In other words, the result of this process was a sequence of party sizes with a fixed mean and standard deviation (across all values), arranged so as to maintain a specified temporal autocorrelation (between adjacent values). The resulting party sizes were then rounded to the nearest integer, with a lower bound of one and an upper bound set by the total group size. 1) With this approach, we aimed to mimic natural *Pan* temporal fluctuations in party size - for example, changes that may arise due to seasonal shifts or other socio-ecological constraints - while retaining control over the overall mean size, variability, and temporal smoothness of party membership.

***2.3 Model optimization***

Although the procedure of resampling individuals into new successive parties involves multiple sequential decisions, in practice we implement all steps using matrix-based operations. By storing initialized dyadic social preferences and party sizes in matrices, and using the Gumbel-max Trick (Huijben et al., 2023) to perform efficient, vectorized softmax operations over arrays, the majority of the simulation can be carried out using linear algebra operations (e.g. matrix multiplication), yielding a scalable method for generating fission-fusion association data.

***2.4 Experimental Design and parameterisation***

**Table S1.** *Descriptions of parameters used to initialize simulations*

| ***Parameter name*** | ***Description*** | ***Used to define:*** | ***Value range*** | ***Sampling increments*** |
| --- | --- | --- | --- | --- |
| *meanf =*  mean difference | Sets the difference between peaks of two normal distributions | Dyadic association preference matrix | 0-100 | 10 |
| *p =*  mixing parameter | The probability of drawing from one of the two normal distributions | Dyadic association preference matrix | 0-0.75 | 0.25 |
| *pref_cv =*  preferences coefficient of variance | The coefficient of variance for each of two normal distributions | Dyadic association preference matrix | 0-0.9 | 0.3 |
| *party_mean =*  mean party size | The mean size of parties for a simulation run | Party size matrix | 5, 9, 13 | NA |
| *party_sd =*  standard deviation of party size | The variance (standard deviation) in party sizes for a simulation run | Party size matrix | 1, 4, 7 | NA |
| *greg_cv =* gregariousness coefficient of variance | Coefficient of variance of normal distribution | List of individual gregariousness | 0.1, 0.4 | NA |

***2.4.1 Using Simple Ration Index for Edge Weights***

We calculated dyadic edge weights for SNA using Simple Ratio Indices (SRI). SRI (Farine, 2013; Whitehead, 1995) is a metric of dyadic association, reflecting the proportion of time two individuals are observed associating relative to the time they are observed but not associating. We calculate SRI as Xab/(Xa+Xb-Xab), where Xab are scans where both individuals a and b were present, Xa are all scans where a was present and Xb are all scans where b was present. In other words, this SRI calculates the proportion of time two individuals were observed associating compared to the total time in which either was observed. We used SRI as this represents a commonly used method for defining edge weights in animal social networks (Farine, 2013; Whitehead, 1995).

***2.4.2 Using STRAND for Edge Weights***

Due to known issues estimating dyadic association given individual differences in sampling, for experiment 2 we additionally estimated dyadic association by fitting STRAND models (Ross et al., 2024), using each individual’s presence in a focal party as the exposure and their co-presence as the outcome. However, under our model where observation at the same time is identical to social association (i.e., unlike grooming, where two individuals can both be observed but not associating with one another, for party association mere co-presence is itself association), mean posterior estimates given by STRAND were nearly identical to those calculated by SRI. We here used flat priors, and the likelihood therefore likely overwhelmed the initial prior and, because co-observation *is* association, is identical to the SRI.

***2.5 Detailed description of network metrics calculated for each simulated network***

1) **Density** (mean edge weight across all dyads). Density represents the average connection strength across the group, essentially measuring the total connectivity across the group (Farine & Whitehead, 2015). Specifically, we calculated density as the proportion of observed ties given possible ties (i.e., the cumulative edge weight divided by the number of possible ties).

2) **Clustering coefficient** (global clustering using Barrat’s method; Barrat et al., 2004). Clustering coefficient represents the degree to which the graph is composed of tightly knit clusters. Specifically, it measures the triangular closure - the weight of closed triplets (sets of three nodes each connected to one another by nonzero edges) to all triplets (sets of three nodes with at least two nonzero edges connecting them). This metric is often used to assess the degree to which “mutual friends” are likely to be close with one another, and by proxy the extent to which the network itself is made up of tightly interconnected clusters.

3) **Modularity** (using Louvain community detection). Modularity represents the degree to which the group is divisible into distinct subgroups. Specifically, it divides the group into cliques using a given community detection algorithm, here using the well-established Louvain algorithm (Blondel et al., 2008), and calculates the relative weight of within- compared to between-clique edges.

4) **Edge CV** (coefficient of variation of *edge weights*). Edge coefficient of variation describes how uneven the pairwise connection strengths are across all dyads in the group. A low edge CV means that most dyads have similar association strength, whereas a high edge CV means that some dyads are strongly connected while others are weakly connected.

5) **Node CV** (coefficient of variation of *node strengths*). Node coefficient of variation describes how uneven the total social connectivity is across individuals. A low node CV means that individuals have a similar overall social connection strength, whereas a high node CV means that some individuals are more strongly connected overall than others.

We plotted all simulated network outputs alongside the same SNA metrics generated from published empirical datasets from bonobos and chimpanzees (ESM 2.6) to assess how our model may relate to real-world systems.

***2.6 Processing published empirical data***

Published datasets contained lists of Simple Ratio Indices for specific date ranges for each species. We combined the published datasets into one data frame so that each dyad was split into two unique individuals. For each dyad we retained information on the community, species (i.e., bonobo, eastern chimpanzees, western chimpanzees), group size (the number of unique individuals included in the dataset), and duration of sampling period (i.e., 3 months, one year). We used this dataset to generate social networks for each community and unique sampling period using the same methodology as for the simulated data and calculated the same SNA metrics. All data processing and network generation can be found on the Github_blinded

***2.7 Experiment parameters***

***2.7.1 Experiment 1***

**Dyadic preference parameters.** Dyadic preferences (entries in the initial symmetric preference matrices) were drawn from a bimodal distribution: each value was sampled from a mixture of two Gaussian distributions with means at *+meanf/2* and *-meanf/2,* with a relative sampling probability of *p* for the first, and 1 - *p* for the second, Gaussian distribution (Figure S1). We swept *meanf* from 0 to 100 in steps of 10, *p* from 0 to 0.75 in steps of 0.25, and the coefficient of variation of preference distributions (*pref_cv*, the same for both distributions) from 0 to 0.9 in steps of 0.3, The *pref_cv* was added to a baseline standard deviation of 0.1 to ensure well-defined distributions even when *meanf* of 0.

**Party parameters.** We varied the average party sizes (*party_mean*) over 5, 9, and 13, and the standard deviation of party sizes (*party_sd*) over 1, 4, and 7.

**Focal sampling.** We used equal focal sampling in all runs, that is, each individual was selected once as focal before any individual was sampled a second time.

***2.7.2 Experiment 2***

**Dyadic preference parameters.** We implemented two main conditions for the preference matrices: 1) without individual gregariousness (i.e., identical to Experiment 1) and 2) with individual gregariousness. For the gregariousness condition, each individual’s gregariousness (i.e., their overall attraction/repulsion to other individuals) was sampled from a normal distribution with mean 0 and coefficient of variation (*greg_cv*) at 0.1 or 0.4. Smaller *greg_cv* indicates more homogeneity of gregariousness across individuals.

In the no-gregariousness condition, we reused the preference matrices from Experiment 1. In the gregariousness condition, we generated new matrices in which composite dyadic preferences were computed as an equal weighting of dyadic preferences (from Experiment 1 matrices - with *pref_p* of 0 [unimodal] or 0.5 [bimodal], with *meanf* fixed at 50 and *pref_cv* at 0.3) and the average gregariousness scores of the two dyad members. In other words, for each dyad of two individuals A and B, and dyadic preference of D_AB (from experiment 1), we assigned individuals gregariousness values of G_A and G_B (generated by the method described above) and calculated the net dyadic preference as D_AB + (G_A + G_B)/2. The composite dyadic preference therefore accounts for each individual’s gregariousness during party resampling, meaning that less gregarious individuals were likely to be observed in parties, even if they shared relatively strong dyadic preferences with individuals in the current party.

**Party parameters.** We fixed *party_mean* at 5 and *party_sd* at 1 to reduce sources of variability to isolate the effects of interest (i.e., individual gregariousness and observation bias). We reused the corresponding party size matrices from Experiment 1.

**Focal sampling.** Individual gregariousness not only influences an individual’s interactions with conspecifics but also the probability of them being seen by human observers. Therefore, variation in individual gregariousness can result in biases during data collection and impact subsequent analyses. We compared three focal sampling schemes: 1) equal sampling (as in Experiment 1, where each individual was sampled once before any were resampled), 2) random sampling (each day, all individuals had equal probability of being selected, with no constraints against repetition), and 3) weighted sampling (used only in the gregariousness condition), in which an individual’s probability of being selected as focal was proportional to their gregariousness score, simulating observer bias toward more gregarious individuals.

***2.7.3 Experiment 3***

**Dyadic preference parameters.** We implemented two main preference conditions: uniform (a matrix of all zeros) and clustered. In the clustered condition, we varied the number of distinct cliques (*n_cliques*). For each run, we first partitioned the group into *n_cliques*, then sampled dyadic preferences from a bimodal distribution as in Experiment 1. However, instead of using *pref_p* to set the mixing proportions, all within-clique dyads were drawn from the higher-mean component, and all between-clique dyads from the lower-mean component. We used *meanf* = 100 and *pref_cv* = 0 (as defined in Experiment 1). As group size increased, we increased the number of possible cliques up to a maximum of 30 (each of size 5 or 6) at the largest group size (155).

**Party parameters.** We again fixed *party_mean* at 5 and *party_sd* at 1.

**Focal sampling.** We used equal focal sampling in all runs.

***2.7.4 Experiment 4***

**Dyadic preference parameters.** We used two conditions (unimodal and bimodal), with a *meanf* fixed at 50 and *pref_cv* at 0.3 for all runs.

**Party size parameters.** We again fixed *party_mean* at 5 and *party_sd* at 1. For the absolute-time condition, we used observation periods of 30, 90, 180, and 360 days. We ran one full simulation for 360 days, and shorter periods were obtained by subsetting the full simulation. For the relative time condition, we used 0.5, 1, 2, and 4 average observation days per individual.

**Focal sampling.** For each parameter combination, we compared equal focal sampling with random focal sampling.

**ESM 3. Correlation between input dyadic association and calculated SRI**

***3.1 Methodology***

For each network, we computed the correlation between measured dyadic SRI values and the dyadic association preferences used to simulate the fission-fusion process to estimate the degree to which the dyadic association patterns computed from the simulations (SRI) capture the pre-defined underlying dyadic association preferences.

***3.2. Results***

Across experiments and simulation runs we found, on average, a positive correlation between the initial dyadic association preferences and dyadic SRI values computed at the end of each simulation run (mean correlation between input preference and SRI across dyads: R^2^ = 0.52, sd = 0.12; Figure S2) indicating that the simulated association data reasonably detected the pre-defined underlying dyadic association preferences. Yet, this relationship varied across runs and between experiments, suggesting that some parameter combinations produced simulated data that more closely reflected the initialized preferences (Figure S2). This variability was largely driven by higher variance in R^2^ for smaller group sizes, likely because the limited set of feasible party sizes in small groups reduces the scope for dyads to display sharply contrasting association preferences.

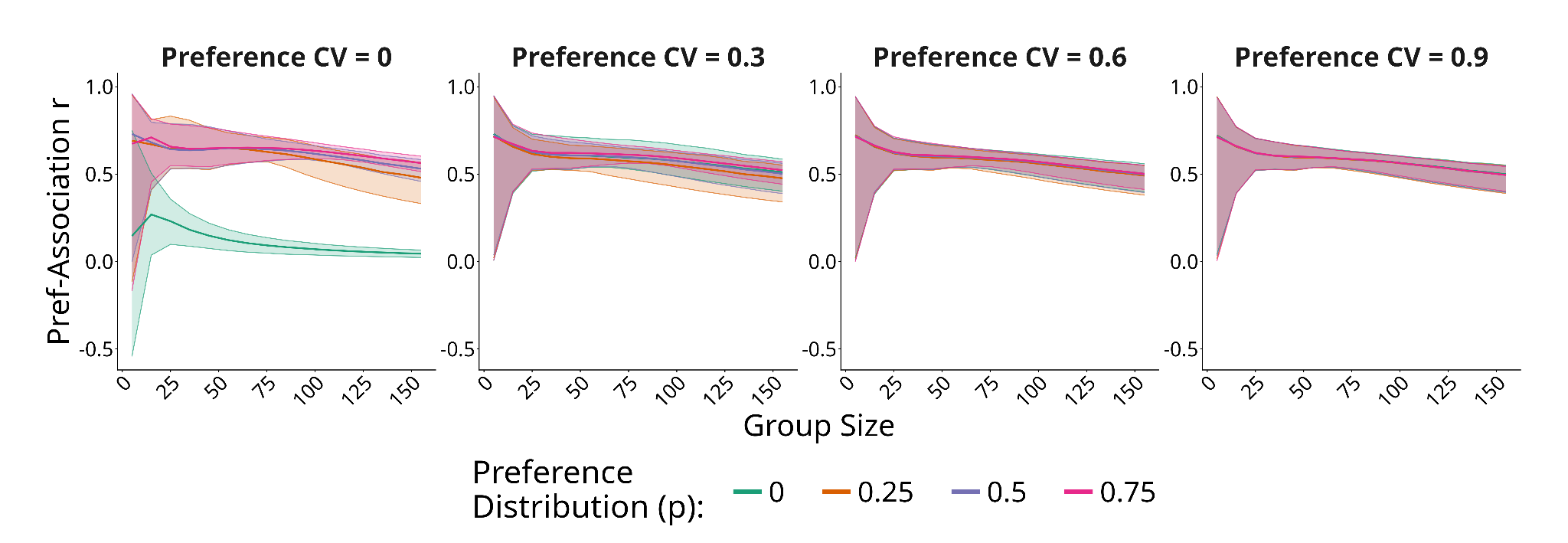

**Figure S2. *Correlation between initial dyadic association preferences and dyadic SRI computed at the end of each simulation in Experiment 1.*** *All correlation across experiments can be found in Github_blinded.*

**ESM 4. Density-relative calculations**

Because it is well-known that many network properties are directly influenced by the density of the network (Faust, 2006), we additionally compared whether estimating global network metrics relative to density (instead of as a raw value) would yield simpler relationships between group size and network metrics.

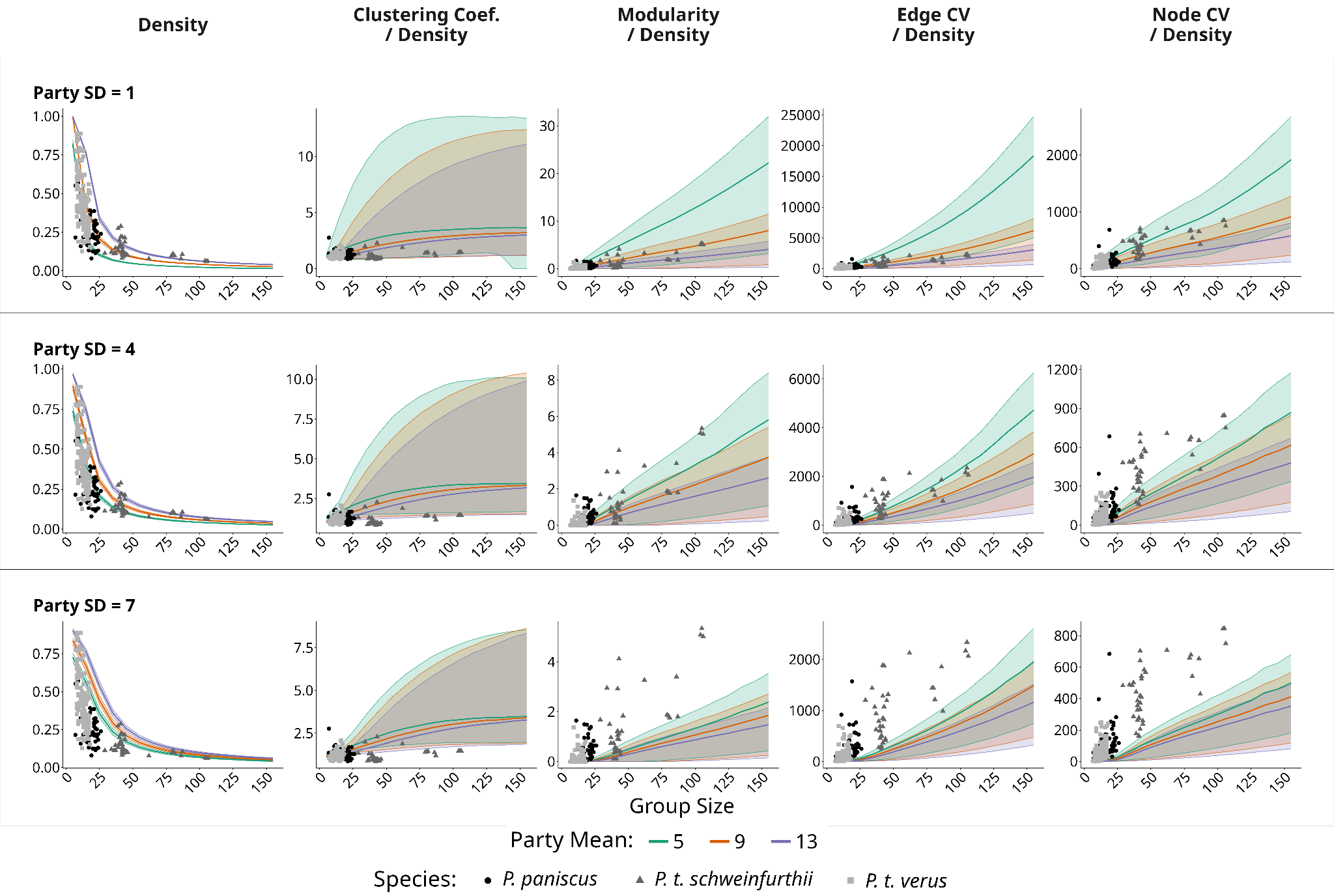

**Figure S3. *Sensitivity of social network metrics, relative to density, across different group and party sizes in experiment 1.*** Changes in social network metrics calculated across simulated runs with varying mean party size (Party Mean), party size variability (Party Standard Deviation: top = low variability, bottom = high variability) and groups of different sizes. Each parameter combination was run 100 times resulting in a total of 2 534 400 simulation runs. Points represent SNA metrics calculated from published empirical data (Samuni et al., 2022; Surbeck et al., 2017).

With this approach, we find that global metrics are relatively “flatter” than the raw values (i.e., main text Figure 2), but neither the empirical nor simulated metrics are constant across changes in, nor follow a simple linear relationship with, group size. Therefore, the issues to comparative inference discussed in the main text focusing on raw metric values cannot be solved simply by taking these values as relative to the overall network density.

**ESM 5. STRAND comparison**

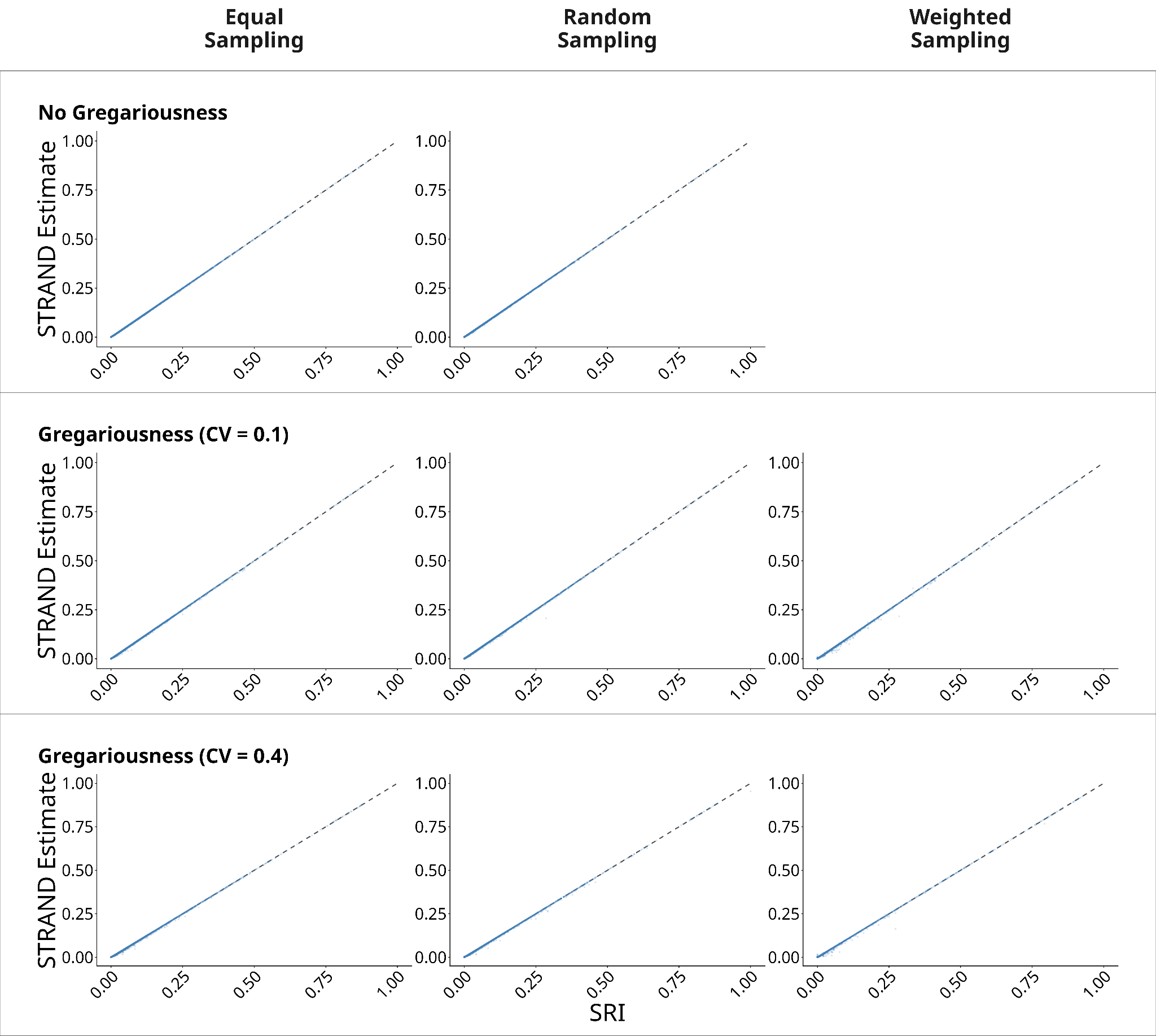

**Figure S4. *Correlation between dyadic SRI and mean of posterior edge weight calculated with STRAND.*** All estimates, across conditions, were nearly identical to the SRI calculation.

**ESM 6. Quantitative difference between empirical and simulated datasets**

***6.1 Experiment 1***

**Table S2.** *Difference between SNA metrics calculated from simulated data and compared to empirical data of the same group size across different mean party sizes of the simulated data. Difference is measured as the value of the simulated data - value from empirical data*

| **Mean party size** | **metric** | **mean** | **sd** | **min** | **max** | **range** |
| --- | --- | --- | --- | --- | --- | --- |
| 5 | density | 0.03 | 0.21 | -0.74 | 0.67 | 1.4 |
| 5 | clustering_coefficient | 0.19 | 0.28 | -0.75 | 0.86 | 1.61 |
| 5 | modularity | 0.01 | 0.11 | -0.39 | 0.48 | 0.87 |
| 5 | cv_edge | -17.28 | 44.97 | -171.07 | 174.56 | 345.62 |
| 5 | cv_node | -14.79 | 16.03 | -70.28 | 54.24 | 124.52 |
| 9 | density | 0.17 | 0.2 | -0.51 | 0.81 | 1.32 |
| 9 | clustering_coefficient | 0.27 | 0.24 | -0.53 | 0.87 | 1.39 |
| 9 | modularity | -0.03 | 0.08 | -0.39 | 0.39 | 0.78 |
| 9 | cv_edge | -30.11 | 34.27 | -172.99 | 78.52 | 251.51 |
| 9 | cv_node | -16.28 | 15.57 | -71.25 | 43.69 | 114.94 |
| 13 | density | 0.3 | 0.2 | -0.18 | 0.81 | 0.99 |
| 13 | clustering_coefficient | 0.37 | 0.21 | -0.16 | 0.87 | 1.03 |
| 13 | modularity | -0.04 | 0.07 | -0.39 | 0.35 | 0.74 |
| 13 | cv_edge | -39.58 | 29.57 | -174.47 | 56.53 | 230.99 |
| 13 | cv_node | -19.11 | 13.69 | -71.76 | 38.79 | 110.54 |

**Table S3.** *Difference between SNA metrics calculated from simulated data and compared to empirical data of the same group size across different levels of variation in sizes of parties of the simulated data. Difference is measured as the value of the simulated data - value from empirical data*

| **SD party sizes** | **metric** | **mean** | **sd** | **min** | **max** | **range** |
| --- | --- | --- | --- | --- | --- | --- |
| 1 | density | 0.11 | 0.27 | -0.74 | 0.81 | 1.55 |
| 1 | clustering_coefficient | 0.15 | 0.29 | -0.75 | 0.87 | 1.62 |
| 1 | modularity | 0.03 | 0.11 | -0.39 | 0.48 | 0.87 |
| 1 | cv_edge | -7.82 | 43.79 | -172.22 | 174.56 | 346.78 |
| 1 | cv_node | -10.8 | 16.84 | -71.76 | 54.24 | 126 |
| 4 | density | 0.17 | 0.21 | -0.54 | 0.79 | 1.33 |
| 4 | clustering_coefficient | 0.3 | 0.21 | -0.41 | 0.87 | 1.28 |
| 4 | modularity | -0.04 | 0.07 | -0.39 | 0.24 | 0.63 |
| 4 | cv_edge | -34.66 | 28.87 | -173.15 | 32.47 | 205.61 |
| 4 | cv_node | -17.96 | 13.5 | -71.51 | 26.18 | 97.69 |
| 7 | density | 0.23 | 0.18 | -0.37 | 0.73 | 1.1 |
| 7 | clustering_coefficient | 0.39 | 0.19 | -0.19 | 0.87 | 1.06 |
| 7 | modularity | -0.06 | 0.07 | -0.39 | 0.17 | 0.56 |
| 7 | cv_edge | -44.47 | 29.17 | -174.47 | 9.73 | 184.19 |
| 7 | cv_node | -21.42 | 13.11 | -71.36 | 9.33 | 80.7 |

**Table S4.** *Difference between SNA metrics calculated from simulated data and compared to empirical data of the same group size across group sizes. Difference is measured as the value of the simulated data - value from empirical data*

| **Group size** | **metric** | **mean** | **sd** | **min** | **max** | **range** |
| --- | --- | --- | --- | --- | --- | --- |
| 5 | density | 0.3 | 0.21 | -0.2 | 0.81 | 1.01 |
| 5 | clustering_coefficient | 0.37 | 0.18 | -0.04 | 0.78 | 0.83 |
| 5 | modularity | -0.02 | 0.05 | -0.26 | 0 | 0.26 |
| 5 | cv_edge | -30.2 | 25.47 | -131.19 | 6.33 | 137.52 |
| 5 | cv_node | -13.81 | 12.58 | -55.37 | 5.65 | 61.02 |
| 15 | density | 0.18 | 0.25 | -0.74 | 0.71 | 1.45 |
| 15 | clustering_coefficient | 0.29 | 0.28 | -0.75 | 0.87 | 1.62 |
| 15 | modularity | -0.02 | 0.09 | -0.26 | 0.44 | 0.71 |
| 15 | cv_edge | -24.3 | 35.41 | -144.97 | 113.23 | 258.21 |
| 15 | cv_node | -13.82 | 14.75 | -62.62 | 54.24 | 116.86 |
| 25 | density | 0.08 | 0.15 | -0.3 | 0.47 | 0.77 |
| 25 | clustering_coefficient | 0.24 | 0.24 | -0.3 | 0.81 | 1.12 |
| 25 | modularity | -0.03 | 0.11 | -0.28 | 0.47 | 0.75 |
| 25 | cv_edge | -24.06 | 39.92 | -134.59 | 116.08 | 250.67 |
| 25 | cv_node | -12.35 | 13.53 | -54.1 | 42.19 | 96.29 |
| 35 | density | 0.05 | 0.1 | -0.22 | 0.28 | 0.5 |
| 35 | clustering_coefficient | 0.21 | 0.23 | -0.26 | 0.83 | 1.09 |
| 35 | modularity | 0 | 0.13 | -0.36 | 0.45 | 0.81 |
| 35 | cv_edge | -48 | 44.97 | -171 | 102.8 | 273.8 |
| 35 | cv_node | -33.38 | 10.5 | -57.93 | 8.07 | 66 |
| 45 | density | 0 | 0.08 | -0.24 | 0.21 | 0.45 |
| 45 | clustering_coefficient | 0.17 | 0.21 | -0.24 | 0.81 | 1.05 |
| 45 | modularity | -0.01 | 0.12 | -0.39 | 0.48 | 0.87 |
| 45 | cv_edge | -40.37 | 49.71 | -174.47 | 140.26 | 314.73 |
| 45 | cv_node | -29.56 | 11.42 | -63.34 | 14.32 | 77.66 |
| 65 | density | 0.02 | 0.04 | -0.05 | 0.1 | 0.15 |
| 65 | clustering_coefficient | 0.1 | 0.19 | -0.12 | 0.68 | 0.79 |
| 65 | modularity | -0.12 | 0.09 | -0.25 | 0.23 | 0.48 |
| 65 | cv_edge | -86 | 46.08 | -158.22 | 73.07 | 231.29 |
| 65 | cv_node | -32.85 | 8.39 | -52.94 | -13.34 | 39.6 |
| 75 | density | -0.02 | 0.03 | -0.08 | 0.05 | 0.13 |
| 75 | clustering_coefficient | 0.08 | 0.18 | -0.13 | 0.65 | 0.78 |
| 75 | modularity | -0.05 | 0.09 | -0.2 | 0.3 | 0.5 |
| 75 | cv_edge | -65.13 | 49.3 | -148.15 | 108.4 | 256.55 |
| 75 | cv_node | -47.24 | 9.31 | -71.76 | -24.64 | 47.12 |
| 85 | density | -0.02 | 0.03 | -0.08 | 0.05 | 0.14 |
| 85 | clustering_coefficient | 0.08 | 0.17 | -0.12 | 0.65 | 0.77 |
| 85 | modularity | -0.07 | 0.1 | -0.26 | 0.3 | 0.56 |
| 85 | cv_edge | -34.33 | 53.45 | -133.02 | 174.56 | 307.58 |
| 85 | cv_node | -28.01 | 8.6 | -51.63 | -7.11 | 44.52 |
| 105 | density | 0 | 0.02 | -0.04 | 0.04 | 0.09 |
| 105 | clustering_coefficient | 0.09 | 0.15 | -0.09 | 0.6 | 0.7 |
| 105 | modularity | -0.18 | 0.09 | -0.32 | 0.17 | 0.5 |
| 105 | cv_edge | -38.51 | 57.04 | -128.36 | 158.53 | 286.89 |
| 105 | cv_node | -28.71 | 8.12 | -51.29 | -8.1 | 43.18 |

***6.2 Experiment 2***

**Table S5.** *Difference between SNA metrics calculated from simulated data and compared to empirical data of the same group size with or without heterogeneity in gregariousness in the simulated data. Difference is measured as the value of the simulated data - value from empirical data*

| **Gregarious condition** | **metric** | **mean** | **sd** | **min** | **max** | **range** |
| --- | --- | --- | --- | --- | --- | --- |
| greg | density | -0.08 | 0.23 | -0.78 | 0.66 | 1.44 |
| greg | clustering_coefficient | -0.1 | 0.27 | -0.89 | 0.74 | 1.63 |
| greg | modularity | 0.13 | 0.12 | -0.37 | 0.54 | 0.91 |
| greg | cv_edge | 56.67 | 93.29 | -128.5 | 1404.13 | 1532.64 |
| greg | cv_node | 9.44 | 30.42 | -54.34 | 313.65 | 367.99 |
| no_greg | density | -0.08 | 0.22 | -0.73 | 0.66 | 1.39 |
| no_greg | clustering_coefficient | -0.09 | 0.26 | -0.74 | 0.74 | 1.48 |
| no_greg | modularity | 0.14 | 0.11 | -0.26 | 0.43 | 0.69 |
| no_greg | cv_edge | 33.86 | 41.28 | -128.02 | 162.53 | 290.55 |
| no_greg | cv_node | -4.82 | 15.97 | -53.6 | 45.9 | 99.5 |

**Table S6.** *Difference between SNA metrics calculated from simulated data and compared to empirical data of the same group size with differences in degree of heterogeneity (coefficient of variance) in gregariousness in the simulated data. Difference is measured as the value of the simulated data - value from empirical data*

| **Gregarious cv** | **metric** | **mean** | **sd** | **min** | **max** | **range** |
| --- | --- | --- | --- | --- | --- | --- |
| 0 | density | -0.08 | 0.22 | -0.73 | 0.66 | 1.39 |
| 0 | clustering_coefficient | -0.09 | 0.26 | -0.74 | 0.74 | 1.48 |
| 0 | modularity | 0.14 | 0.11 | -0.26 | 0.43 | 0.69 |
| 0 | cv_edge | 33.86 | 41.28 | -128.02 | 162.53 | 290.55 |
| 0 | cv_node | -4.82 | 15.97 | -53.6 | 45.9 | 99.5 |
| 0.1 | density | -0.08 | 0.23 | -0.74 | 0.66 | 1.41 |
| 0.1 | clustering_coefficient | -0.1 | 0.26 | -0.76 | 0.74 | 1.5 |
| 0.1 | modularity | 0.14 | 0.12 | -0.26 | 0.54 | 0.79 |
| 0.1 | cv_edge | 44.47 | 62.92 | -128.12 | 846.46 | 974.58 |
| 0.1 | cv_node | 1.39 | 19.83 | -54.12 | 207.68 | 261.8 |
| 0.4 | density | -0.09 | 0.23 | -0.78 | 0.66 | 1.44 |
| 0.4 | clustering_coefficient | -0.11 | 0.27 | -0.89 | 0.74 | 1.63 |
| 0.4 | modularity | 0.11 | 0.11 | -0.37 | 0.45 | 0.82 |
| 0.4 | cv_edge | 68.87 | 114.67 | -128.5 | 1404.13 | 1532.64 |
| 0.4 | cv_node | 17.49 | 36.45 | -54.34 | 313.65 | 367.99 |

**Table S7.** *Difference between SNA metrics calculated from simulated data and compared to empirical data of the same group size with different focal sampling methods used to generate the simulated data. Difference is measured as the value of the simulated data - value from empirical data*

| **Focal method** | **metric** | **mean** | **sd** | **min** | **max** | **range** |
| --- | --- | --- | --- | --- | --- | --- |
| equal | density | -0.08 | 0.22 | -0.73 | 0.66 | 1.39 |
| equal | clustering_coefficient | -0.09 | 0.26 | -0.74 | 0.74 | 1.48 |
| equal | modularity | 0.14 | 0.11 | -0.26 | 0.41 | 0.67 |
| equal | cv_edge | 36.71 | 48.48 | -127.62 | 538.86 | 666.48 |
| equal | cv_node | -1.69 | 16.94 | -53.53 | 142.46 | 195.98 |
| random | density | -0.08 | 0.22 | -0.73 | 0.66 | 1.39 |
| random | clustering_coefficient | -0.09 | 0.26 | -0.74 | 0.74 | 1.49 |
| random | modularity | 0.14 | 0.11 | -0.26 | 0.43 | 0.69 |
| random | cv_edge | 38.47 | 50.97 | -128.02 | 563.36 | 691.38 |
| random | cv_node | -0.9 | 17.17 | -53.64 | 143.59 | 197.23 |
| weighted | density | -0.1 | 0.23 | -0.78 | 0.66 | 1.44 |
| weighted | clustering_coefficient | -0.13 | 0.28 | -0.89 | 0.74 | 1.63 |
| weighted | modularity | 0.12 | 0.13 | -0.37 | 0.54 | 0.91 |
| weighted | cv_edge | 91.1 | 136.49 | -128.5 | 1404.13 | 1532.64 |
| weighted | cv_node | 27.39 | 41.17 | -54.34 | 313.65 | 367.99 |

***7.3 Experiment 3***

**Table S8.** *Difference between SNA metrics calculated from simulated data and compared to empirical data of the same group size across clustered or uniform conditions used to generate the simulated data. Difference is measured as the value of the simulated data - value from empirical data.*

| **Cluster consition** | **metric** | **mean** | **sd** | **min** | **max** | **range** |
| --- | --- | --- | --- | --- | --- | --- |
| clustered | density | -0.08 | 0.17 | -0.71 | 0.66 | 1.38 |
| clustered | clustering_coefficient | -0.05 | 0.22 | -0.76 | 0.73 | 1.49 |
| clustered | modularity | 0.39 | 0.23 | -0.26 | 0.79 | 1.05 |
| clustered | cv_edge | 88.54 | 74.76 | -127.17 | 289.2 | 416.37 |
| clustered | cv_node | -18.26 | 16.52 | -67.08 | 30.51 | 97.59 |
| uniform | density | -0.08 | 0.22 | -0.71 | 0.66 | 1.38 |
| uniform | clustering_coefficient | 0.4 | 0.16 | -0.1 | 0.78 | 0.88 |
| uniform | modularity | -0.07 | 0.07 | -0.39 | 0 | 0.39 |
| uniform | cv_edge | -48.04 | 31.43 | -158.05 | 2.3 | 160.35 |
| uniform | cv_node | -24.57 | 15.52 | -68.99 | 0.68 | 69.68 |

**Table S9.** *Difference between SNA metrics calculated from simulated data and compared to empirical data of the same group size across different numbers of cliques in the simulated data. Difference is measured as the value of the simulated data - value from empirical data*

| **Number cliques** | **metric** | **mean** | **sd** | **min** | **max** | **range** |
| --- | --- | --- | --- | --- | --- | --- |
| 1 | density | -0.08 | 0.22 | -0.71 | 0.66 | 1.38 |
| 1 | clustering_coefficient | 0.18 | 0.3 | -0.69 | 0.78 | 1.47 |
| 1 | modularity | 0 | 0.11 | -0.39 | 0.29 | 0.67 |
| 1 | cv_edge | -17.03 | 45.07 | -158.05 | 111.27 | 269.32 |
| 1 | cv_node | -18 | 16.69 | -68.99 | 26.57 | 95.56 |
| 2 | density | -0.05 | 0.21 | -0.67 | 0.66 | 1.34 |
| 2 | clustering_coefficient | -0.06 | 0.26 | -0.75 | 0.73 | 1.48 |
| 2 | modularity | 0.32 | 0.19 | -0.26 | 0.53 | 0.78 |
| 2 | cv_edge | 53.08 | 47.99 | -126.02 | 157.55 | 283.57 |
| 2 | cv_node | -7.61 | 13.38 | -52.76 | 30.51 | 83.27 |
| 3 | density | -0.12 | 0.14 | -0.66 | 0.17 | 0.83 |
| 3 | clustering_coefficient | -0.17 | 0.22 | -0.76 | 0.29 | 1.05 |
| 3 | modularity | 0.47 | 0.08 | 0.2 | 0.65 | 0.45 |
| 3 | cv_edge | 86.09 | 29.22 | -28.66 | 182.63 | 211.29 |
| 3 | cv_node | -15.92 | 11.58 | -59.47 | 18.51 | 77.98 |
| 4 | density | -0.08 | 0.06 | -0.25 | 0.04 | 0.29 |
| 4 | clustering_coefficient | 0.02 | 0.17 | -0.31 | 0.37 | 0.68 |
| 4 | modularity | 0.59 | 0.08 | 0.35 | 0.72 | 0.37 |
| 4 | cv_edge | 118.87 | 29.52 | 28.48 | 198.61 | 170.13 |
| 4 | cv_node | -20.4 | 8.76 | -47.9 | 5.29 | 53.19 |
| 5 | density | -0.08 | 0.06 | -0.25 | 0.03 | 0.28 |
| 5 | clustering_coefficient | 0 | 0.19 | -0.31 | 0.4 | 0.71 |
| 5 | modularity | 0.58 | 0.11 | 0.31 | 0.77 | 0.46 |
| 5 | cv_edge | 126.56 | 32.31 | 35.3 | 215.81 | 180.52 |
| 5 | cv_node | -25 | 10.29 | -49.78 | -2.38 | 47.4 |
| 6 | density | -0.07 | 0.05 | -0.22 | -0.01 | 0.21 |
| 6 | clustering_coefficient | 0.02 | 0.1 | -0.26 | 0.23 | 0.49 |
| 6 | modularity | 0.65 | 0.09 | 0.37 | 0.79 | 0.43 |
| 6 | cv_edge | 145.72 | 35.65 | 46.29 | 226.18 | 179.89 |
| 6 | cv_node | -32.5 | 11.39 | -52.02 | -2.5 | 49.52 |
| 7 | density | -0.07 | 0.05 | -0.22 | 0 | 0.21 |
| 7 | clustering_coefficient | -0.02 | 0.11 | -0.27 | 0.27 | 0.54 |
| 7 | modularity | 0.62 | 0.1 | 0.27 | 0.78 | 0.51 |
| 7 | cv_edge | 148.76 | 40.41 | 35.66 | 238.27 | 202.61 |
| 7 | cv_node | -36.23 | 11.69 | -55.51 | -2.51 | 53 |
| 8 | density | -0.07 | 0.05 | -0.22 | -0.01 | 0.21 |
| 8 | clustering_coefficient | 0.01 | 0.14 | -0.24 | 0.32 | 0.56 |
| 8 | modularity | 0.61 | 0.07 | 0.33 | 0.74 | 0.41 |
| 8 | cv_edge | 164.7 | 39.5 | 68.67 | 245.9 | 177.23 |
| 8 | cv_node | -35.43 | 12.49 | -57.96 | -4.21 | 53.75 |
| 9 | density | -0.07 | 0.05 | -0.22 | -0.01 | 0.21 |
| 9 | clustering_coefficient | 0.01 | 0.15 | -0.24 | 0.33 | 0.57 |
| 9 | modularity | 0.57 | 0.09 | 0.25 | 0.71 | 0.47 |
| 9 | cv_edge | 163.25 | 45.15 | 57.77 | 265.14 | 207.37 |
| 9 | cv_node | -37.27 | 12.48 | -59.1 | -4.96 | 54.14 |
| 10 | density | -0.05 | 0.02 | -0.07 | -0.02 | 0.05 |
| 10 | clustering_coefficient | 0.19 | 0.09 | -0.09 | 0.35 | 0.45 |
| 10 | modularity | 0.64 | 0.06 | 0.56 | 0.73 | 0.16 |
| 10 | cv_edge | 216.19 | 33.68 | 141.41 | 264.75 | 123.34 |
| 10 | cv_node | -28.99 | 13.18 | -56.66 | -6.32 | 50.34 |
| 11 | density | -0.05 | 0.01 | -0.07 | -0.02 | 0.05 |
| 11 | clustering_coefficient | 0.15 | 0.12 | -0.09 | 0.31 | 0.41 |
| 11 | modularity | 0.63 | 0.06 | 0.53 | 0.73 | 0.2 |
| 11 | cv_edge | 221.33 | 36.86 | 139.89 | 271.87 | 131.98 |
| 11 | cv_node | -32.29 | 13.9 | -61.46 | -10.32 | 51.13 |
| 12 | density | -0.05 | 0.01 | -0.07 | -0.03 | 0.04 |
| 12 | clustering_coefficient | 0.12 | 0.15 | -0.1 | 0.35 | 0.45 |
| 12 | modularity | 0.62 | 0.06 | 0.48 | 0.73 | 0.25 |
| 12 | cv_edge | 224.39 | 39.54 | 133.17 | 275.57 | 142.4 |
| 12 | cv_node | -35.6 | 13.44 | -63.29 | -14.9 | 48.38 |
| 13 | density | -0.05 | 0.01 | -0.07 | -0.03 | 0.04 |
| 13 | clustering_coefficient | 0.08 | 0.16 | -0.11 | 0.38 | 0.49 |
| 13 | modularity | 0.6 | 0.06 | 0.43 | 0.7 | 0.28 |
| 13 | cv_edge | 224.7 | 43.37 | 127.55 | 278.82 | 151.28 |
| 13 | cv_node | -38.23 | 12.56 | -65.03 | -18.1 | 46.93 |
| 14 | density | -0.05 | 0.02 | -0.07 | -0.03 | 0.04 |
| 14 | clustering_coefficient | 0.05 | 0.13 | -0.1 | 0.29 | 0.4 |
| 14 | modularity | 0.59 | 0.03 | 0.54 | 0.67 | 0.13 |
| 14 | cv_edge | 236.2 | 34.74 | 168.4 | 284.5 | 116.1 |
| 14 | cv_node | -39.84 | 12.13 | -65.99 | -21.67 | 44.32 |
| 15 | density | -0.05 | 0.02 | -0.07 | -0.03 | 0.04 |
| 15 | clustering_coefficient | 0.02 | 0.11 | -0.11 | 0.22 | 0.33 |
| 15 | modularity | 0.57 | 0.03 | 0.49 | 0.65 | 0.16 |
| 15 | cv_edge | 234.51 | 37.85 | 160.48 | 289.2 | 128.72 |
| 15 | cv_node | -41.78 | 11.2 | -67.08 | -26.17 | 40.91 |
| 16 | density | -0.04 | 0.02 | -0.07 | -0.03 | 0.04 |
| 16 | clustering_coefficient | 0.01 | 0.07 | -0.1 | 0.13 | 0.23 |
| 16 | modularity | 0.55 | 0.03 | 0.48 | 0.62 | 0.14 |
| 16 | cv_edge | 251.32 | 23.21 | 199.73 | 288.64 | 88.91 |
| 16 | cv_node | -37.34 | 4.06 | -46.96 | -28.01 | 18.95 |
| 17 | density | -0.04 | 0.02 | -0.07 | -0.03 | 0.04 |
| 17 | clustering_coefficient | -0.02 | 0.06 | -0.1 | 0.09 | 0.19 |
| 17 | modularity | 0.53 | 0.03 | 0.44 | 0.59 | 0.15 |
| 17 | cv_edge | 248.7 | 25.21 | 194.16 | 289.09 | 94.93 |
| 17 | cv_node | -38.82 | 3.68 | -46.53 | -29.71 | 16.82 |
| 18 | density | -0.03 | 0 | -0.03 | -0.03 | 0.01 |
| 18 | clustering_coefficient | 0.01 | 0.01 | -0.02 | 0.03 | 0.05 |
| 18 | modularity | 0.51 | 0.01 | 0.47 | 0.54 | 0.06 |
| 18 | cv_edge | 268.97 | 6.76 | 249.26 | 289.12 | 39.86 |
| 18 | cv_node | -38.49 | 3.13 | -44.78 | -31.95 | 12.83 |
| 19 | density | -0.03 | 0 | -0.03 | -0.03 | 0.01 |
| 19 | clustering_coefficient | -0.01 | 0.01 | -0.04 | 0.01 | 0.05 |
| 19 | modularity | 0.48 | 0.01 | 0.45 | 0.52 | 0.06 |
| 19 | cv_edge | 264.89 | 6.87 | 248.07 | 284.23 | 36.16 |
| 19 | cv_node | -39.19 | 3.12 | -44.61 | -32.59 | 12.02 |
| 20 | density | -0.03 | 0 | -0.03 | -0.03 | 0.01 |
| 20 | clustering_coefficient | -0.02 | 0.01 | -0.04 | 0.01 | 0.05 |
| 20 | modularity | 0.46 | 0.01 | 0.43 | 0.49 | 0.06 |
| 20 | cv_edge | 261.47 | 6.98 | 245.51 | 281.01 | 35.5 |
| 20 | cv_node | -39.45 | 3.16 | -45.18 | -31.58 | 13.61 |
| 21 | density | -0.03 | 0 | -0.04 | -0.03 | 0.01 |
| 21 | clustering_coefficient | -0.03 | 0.01 | -0.05 | -0.01 | 0.04 |
| 21 | modularity | 0.43 | 0.01 | 0.41 | 0.47 | 0.06 |
| 21 | cv_edge | 257.3 | 7.26 | 239.92 | 279.1 | 39.18 |
| 21 | cv_node | -40.09 | 3.18 | -45.74 | -32.82 | 12.92 |

***6.4 Experiment 4***

**Table S10.** *Difference between SNA metrics calculated from simulated data and compared to empirical data of the same group size across different numbers of absolute days used to generate the simulated data. Difference is measured as the value of the simulated data - value from empirical data*

| **Number days** | **metric** | **mean** | **sd** | **min** | **max** | **range** |
| --- | --- | --- | --- | --- | --- | --- |
| 30 | density | -0.08 | 0.22 | -0.73 | 0.69 | 1.42 |
| 30 | clustering_coefficient | -0.09 | 0.26 | -0.75 | 0.75 | 1.5 |
| 30 | modularity | 0.16 | 0.12 | -0.26 | 0.48 | 0.74 |
| 30 | cv_edge | 41.72 | 49.06 | -127.86 | 261.4 | 389.26 |
| 30 | cv_node | -1.83 | 15.3 | -53.98 | 48.89 | 102.87 |
| 90 | density | -0.08 | 0.22 | -0.73 | 0.67 | 1.39 |
| 90 | clustering_coefficient | -0.09 | 0.26 | -0.74 | 0.73 | 1.47 |
| 90 | modularity | 0.14 | 0.11 | -0.26 | 0.45 | 0.7 |
| 90 | cv_edge | 34.28 | 41.57 | -127.85 | 166.02 | 293.87 |
| 90 | cv_node | -4.49 | 15.89 | -53.59 | 44.78 | 98.37 |
| 180 | density | -0.08 | 0.22 | -0.72 | 0.66 | 1.38 |
| 180 | clustering_coefficient | -0.09 | 0.26 | -0.74 | 0.75 | 1.48 |
| 180 | modularity | 0.14 | 0.11 | -0.26 | 0.43 | 0.68 |
| 180 | cv_edge | 32.5 | 40.15 | -127.73 | 142 | 269.73 |
| 180 | cv_node | -5.15 | 16.13 | -53.88 | 44.12 | 98 |
| 360 | density | -0.08 | 0.22 | -0.72 | 0.65 | 1.37 |
| 360 | clustering_coefficient | -0.09 | 0.26 | -0.73 | 0.75 | 1.48 |
| 360 | modularity | 0.14 | 0.11 | -0.26 | 0.43 | 0.69 |
| 360 | cv_edge | 31.55 | 39.53 | -127.39 | 128.88 | 256.27 |
| 360 | cv_node | -5.46 | 16.25 | -54.29 | 42.98 | 97.26 |

**Table S11.** *Difference between SNA metrics calculated from simulated data and compared to empirical data of the same group size across different numbers of focal days per individual used to generate the simulated data. Difference is measured as the value of the simulated data - value from empirical data*

| **Number days** | **metric** | **mean** | **sd** | **min** | **max** | **range** |
| --- | --- | --- | --- | --- | --- | --- |
| 0.5 | density | -0.08 | 0.23 | -0.76 | 0.78 | 1.54 |
| 0.5 | clustering_coefficient | -0.1 | 0.25 | -0.89 | 0.75 | 1.64 |
| 0.5 | modularity | 0.16 | 0.13 | -0.26 | 0.5 | 0.76 |
| 0.5 | cv_edge | 51.14 | 48.48 | -129.69 | 202.97 | 332.66 |
| 0.5 | cv_node | 3.95 | 18.57 | -54.65 | 80.74 | 135.39 |
| 1 | density | -0.08 | 0.22 | -0.74 | 0.75 | 1.49 |
| 1 | clustering_coefficient | -0.09 | 0.25 | -0.89 | 0.76 | 1.64 |
| 1 | modularity | 0.15 | 0.12 | -0.26 | 0.45 | 0.71 |
| 1 | cv_edge | 39.89 | 43.06 | -128.78 | 161.95 | 290.72 |
| 1 | cv_node | -1.57 | 16.98 | -54.37 | 68.72 | 123.09 |
| 2 | density | -0.08 | 0.22 | -0.74 | 0.71 | 1.45 |
| 2 | clustering_coefficient | -0.09 | 0.26 | -0.75 | 0.74 | 1.49 |
| 2 | modularity | 0.14 | 0.12 | -0.26 | 0.42 | 0.68 |
| 2 | cv_edge | 35.16 | 40.83 | -128.31 | 138.53 | 266.84 |
| 2 | cv_node | -3.9 | 16.55 | -54.04 | 63.43 | 117.47 |
| 4 | density | -0.08 | 0.22 | -0.73 | 0.69 | 1.41 |
| 4 | clustering_coefficient | -0.09 | 0.26 | -0.75 | 0.74 | 1.49 |
| 4 | modularity | 0.14 | 0.11 | -0.26 | 0.42 | 0.67 |
| 4 | cv_edge | 32.62 | 39.72 | -128.28 | 130.12 | 258.4 |
| 4 | cv_node | -4.98 | 16.36 | -54.4 | 48.45 | 102.85 |

**References**

Aureli, F., Schaffner, C. M., Boesch, C., Bearder, S. K., Call, J., Chapman, C. A., Connor, R., Fiore, A. D., Dunbar, R. I. M., Henzi, S. P., Holekamp, K., Korstjens, A. H., Layton, R., Lee, P., Lehmann, J., Manson, J. H., Ramos‐Fernandez, G., Strier, K. B., & Schaik, C. P. van. (2008). Fission‐Fusion Dynamics: New Research Frameworks. *Current Anthropology*, *49*(4), 627–654. https://doi.org/10.1086/586708

Barrat, A., Barthélemy, M., Pastor-Satorras, R., & Vespignani, A. (2004). The architecture of complex weighted networks. *Proceedings of the National Academy of Sciences*, *101*(11), 3747–3752. https://doi.org/10.1073/pnas.0400087101

Blondel, V. D., Guillaume, J.-L., Lambiotte, R., & Lefebvre, E. (2008). Fast unfolding of communities in large networks. *Journal of Statistical Mechanics: Theory and Experiment*, *2008*(10), P10008. https://doi.org/10.1088/1742-5468/2008/10/P10008

Farine, D. R., & Whitehead, H. (2015). Constructing, conducting and interpreting animal social network analysis. *Journal of Animal Ecology*, *84*(5), 1144–1163. https://doi.org/10.1111/1365-2656.12418

Faust, K. (2006). Comparing social networks: Size, density, and local structure. *Advances in Methodology and Statistics*, *3*(2). https://doi.org/10.51936/sdbv3216

Huijben, I. A. M., Kool, W., Paulus, M. B., & van Sloun, R. J. G. (2023). A Review of the Gumbel-max Trick and its Extensions for Discrete Stochasticity in Machine Learning. *IEEE Transactions on Pattern Analysis and Machine Intelligence*, *45*(2), 1353–1371. https://doi.org/10.1109/TPAMI.2022.3157042

Kano, T. (1982). The social group of pygmy chimpanzees (Pan paniscus) of Wamba. *Primates*, *23*(2), Article 2. https://doi.org/10.1007/BF02381159

Madsen, A., & De Silva, S. (2024). Societies with fission–fusion dynamics as complex adaptive systems: The importance of scale. *Philosophical Transactions of the Royal Society B: Biological Sciences*, *379*(1909), 20230175. https://doi.org/10.1098/rstb.2023.0175

Nishida, T. (1968). The social group of wild chimpanzees in the Mahale Mountains. *Primates*, *7*, 167–224.

Samuni, L., Langergraber, K. E., & Surbeck, M. H. (2022). Characterization of *Pan* social systems reveals in-group/out-group distinction and out-group tolerance in bonobos. *Proceedings of the National Academy of Sciences*, *119*(26), e2201122119. https://doi.org/10.1073/pnas.2201122119

Surbeck, M., Girard-Buttoz, C., Boesch, C., Crockford, C., Fruth, B., Hohmann, G., Langergraber, K. E., Zuberbühler, K., Wittig, R. M., & Mundry, R. (2017). Sex-specific association patterns in bonobos and chimpanzees reflect species differences in cooperation. *Royal Society Open Science*, *4*(5), Article 5. https://doi.org/10.1098/rsos.161081
